# Cell Division History Determines Hematopoietic Stem Cell Potency

**DOI:** 10.1101/503813

**Authors:** Fumio Arai, Patrick S. Stumpf, Yoshiko M. Ikushima, Kentaro Hosokawa, Aline Roch, Matthias P. Lutolf, Toshio Suda, Ben D. MacArthur

## Abstract

Loss of stem cell self-renewal may underpin aging. Here, we combined single cell profiling, deep-learning, mathematical modelling and *in vivo* functional studies to explore how hematopoietic stem cell (HSC) division patterns evolve with age. We trained an artificial neural network (ANN) to accurately identify cell types in the hematopoietic hierarchy and predict their age from their gene expression patterns. We then used this ANN to compare daughter cell identities immediately after HSC divisions, and found that HSC self-renewal declines sharply with age. Furthermore, while HSC cell divisions are deterministic and intrinsically-regulated in young and old age, they become stochastic and niche-sensitive in mid-life. Analysis of evolving division patterns indicated that the self-renewal ability an individual HSC depends upon number times it has divided previously. We propose a model of HSC proliferation that accurately predicts the accumulation of HSCs with age, and reconciles the stochastic and instructive views of fate commitment.

## INTRODUCTION

Hematopoietic stem cells (HSCs) are able to give rise to all cell types in the hematopoietic lineages, and are responsible for ensuring healthy life-long hematopoiesis (Orkin & Zon 2008). To do so, they must regulate their own numbers and simultaneously produce differentiated progeny in precisely regulated proportions. Maintaining this balance requires HSCs to have the potential to undergo both symmetric and asymmetric cell divisions, and appropriately alter their division patterns in response to environmental cues (Morrison & Kimble 2006).

Asymmetric cell divisions produce two daughter cells of different ‘types’. For stem cells this typically means the production of one self-renewing daughter cell (i.e. another stem cell) and one differentiated daughter cell that does not retain self-renewal potential. By contrast, symmetric stem cell divisions produce two daughter cells of the same type: either two stem cells, or two progenitor cells that have undergone the first step of commitment. Because asymmetric cell divisions produce one stem cell per division, they are a natural way to maintain stem cell numbers while simultaneously producing differentiated progeny, and have accordingly been widely studied as a mechanism for maintaining tissue homeostasis (Yamashita et al. 2010). However, there is growing evidence that tissue homeostasis can also be maintained by balancing different kinds of symmetric divisions within the stem cell pool (Simons & Clevers 2011). In particular, if the propensity for each stem cell to divide symmetrically to produce two committed progenitors is equal to the propensity to symmetrically divide to produce two stem cells then the expected number of stem cells will not change. In this case, the propensities for different kinds of symmetric and asymmetric divisions must be tightly controlled to ensure an appropriate stem cell pool is maintained with age. Failure of this control can have dramatic consequences for the organism. For example, various different types of cancer, including leukemia, can be caused by loss of balance in cell division patterns (Alcolea et al. 2014, Frede et al. 2016, Stiehl & Marciniak-Czochra 2017, Wu et al. 2007, Zimdahl et al. 2014).

To achieve this control, HSC dynamics are regulated by extracellular signals from the specialized bone marrow microenvironment(s), or niche(s), in which they reside (Moore & Lemischka 2006, Wilson & Trumpp 2006). Instruction from the niche appears to be particularly important for maintaining tissue homeostasis by producing a stem cell population that is quiescent under normal circumstances, yet poised to initiate appropriate rapid tissue expansion when needed – for example, on exposure to stress stimuli (Trumpp et al. 2010, Mendelson & Frenette 2014, Prendergast & Essers 2014, Karigane et al. 2016, van Galen et al. 2014).However, stem cell dynamics are not determined by the niche alone, but rather by an interplay between cell intrinsic factors and niche instruction.

For example, in the Drosophila testis, signals from niche hub cells induce asymmetric localization of adenomatous polyposis coli (APC) – a component of the Wnt signalling pathway that regulates proliferation – which results in stem cell divisions that are perpendicular to hub cells. Daughter cells that maintain contact with the hub cells maintain APC expression and retain a stem cell identity, while those that are separated from the hub cell niche undergo differentiation (Yamashita et al. 2003). Similar niche signals, in the form of cell-cell contacts have also been shown to control spindle orientations and induce asymmetric cell divisions in other types of stem and progenitor cells, including Drosophila neural precursor cells and mouse epidermal stem cells (Bhat 2014, Lechler & Fuchs 2005).

These results indicate that niche signals can unambiguously instruct cell division events, and appear to argue against the stochastic view of stem cell proliferation in which different kinds of division occur with defined probabilities. However, it is not clear if niche instruction is always so precise. For some mammalian stem cell types, including HSCs, it has long been suggested that the niche only *probabilistically* exerts its control (Till et al. 1964, Nakahata et al. 1982, Suda et al. 1984). In this view, the niche does not precisely instruct the developmental trajectory of each individual HSC, but rather tunes the probabilities with which different kinds of symmetric and asymmetric divisions will occur within the stem cell pool as a whole. Thus, while the stem cell population may be well-regulated, individual cells may not be closely controlled (Till et al. 1964). This view is consistent with clone size distributions from colony forming assays both *in vivo* and *in vitro* (Till et al. 1964, Clayton et al. 2007, Klein & Simons 2011, Greulich & Simons 2016) and with experiments which show that niche activity is positively associated with HSC numbers (Ding et al. 2012, Zhang et al. 2003, Calvi et al. 2003, Kunisaki et al. 2013, Visnjic et al. 2004, Méndez-Ferrer et al. 2010).

Collectively, these results indicate that both stochastic and deterministic mechanisms have a role in regulating stem cell proliferation. However, it is not clear how these different mechanisms work collectively to guide stem cell dynamics, and how the balance between them changes with age. Here, we sought to use single cell profiling, machine-learning and mathematical modeling to address this question. We find evidence for both stochastic and instructive modes of regulation, yet also observe that the balance between stochastic and instructive mechanisms changes substantially with age. Based on these results we propose a model stem cell proliferation that combines stochastic and niche-instructive processes and accurately predicts how stem cell numbers change with age.

## RESULTS

### The Evi1 positive HSC fraction is enriched for LT-HSCs

To begin our investigation we sought to isolate a highly-purified, homogeneous, population of long-term repopulating HSCs (LT-HSCs). It has previously been shown that ecotropic viral integration site 1 (Evi1) (also known as MDS1 and EVI1 complex locus, or Mecom) is positively associated with HSC activity in the immunophenotypic stem cell pool (Goyama et al. 2008, Kataoka et al. 2011, Yuasa et al. 2005). We therefore utilized an Evi1 green fluorescent protein (GFP) knock-in mouse model (Kataoka et al. 2011) to select Lineage^−^Sca-1^+^c-Kit^+^ (LSK) CD41^−^CD48^−^CD150^+^CD34^+^ (representing a heterogeneous mixture of hematopoietic stem/progenitor cells, denoted CD34^+^), LSKCD41^−^CD48^−^CD150^+^CD34^−^ Evi1-GFP^+^ (denoted Evi1^+^) and LSKCD41^−^CD48^−^CD150^+^CD34^−^ Evi1-GFP^−^ (denoted Evi1^−^) fractions (**Figure 1A**) from the bone marrow (BM) of 8-week old mice.

**Figure 1.**
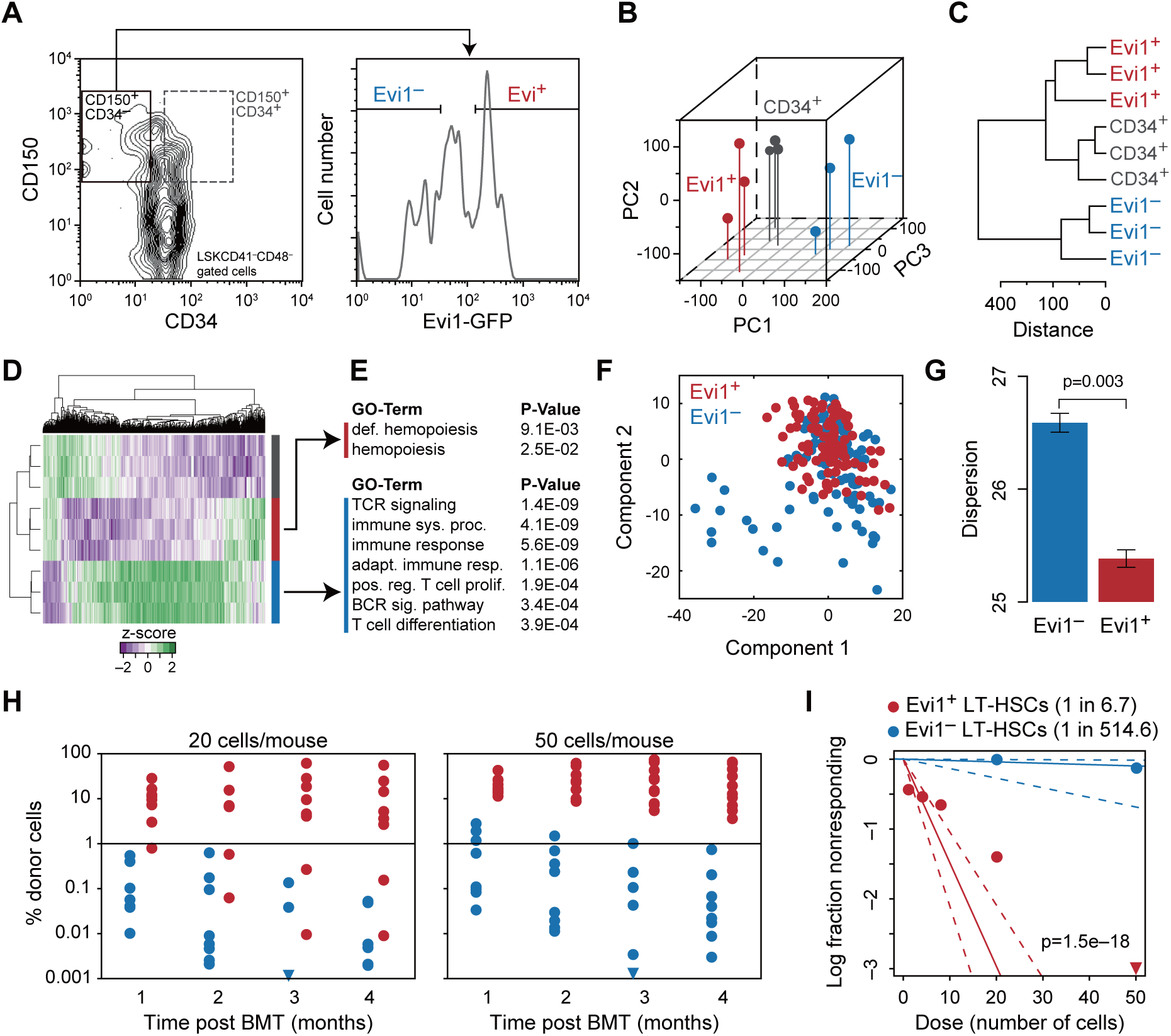
Evi1 expression is positively associated LT-HSC activity. (A) Selection protocol. LSKCD41^−^CD48^−^CD150^+^CD34^+^ (dotted square in panel A, denoted CD34^+^), LSKCD41^−^CD48^−^CD150^+^CD34^−^ Evi1^+^ (denoted Evi1^+^) and LSKCD41^−^CD48^−^CD150^+^CD34^−^ Evi1^−^ (denoted Evi1^−^) fractions were isolated from the BM of 8-week old mice. (B-C) CD150^+^CD34^+^, Evi1^+^ and Evi1^−^ fractions form distinct populations based upon global gene expression patterns. (B) Projection of global expression patterns on to the first three principal components (PCs, three biological replicates per population). (C) Hierarchical clustering of global expression patterns (3 biological replicates per population). (D) Heatmap of global expression patterns shows that each population is characterized by different combinations of gene expression. (E) Genes associated with definitive hemopoiesis are upregulated in the Evi1^+^ fraction; genes associated differentiation are upregulated in the Evi1^−^ fraction. (F-G) The Evi1^+^ is more homogeneous than the Evi1^−^ fraction. (F) Projection of single cell expression patterns on to the first two PCs. (G) Total dispersion in the Evi1^+^ and Evi1^−^ fractions. Data is presented as mean ± standard deviation, p-value is from two sided t-test (96 cells in each group) (H-I) Results of bone marrow transplantations. Evi1^+^ cells show better engraftment as assessed by: (H) percentage donor derived cells in peripheral blood and (I) enrichment for functional LT-HSCs. p-values is from extreme value limiting dilution analysis (ELDA) (Hu & Smyth 2009).

To investigate differences in global patterns of gene expression, we profiled these populations using microarrays. We found that the Evi1^+^ fraction had distinct patterns of gene expression, and could be discriminated from both the CD34^+^ and Evi1^−^ fractions, based upon these differences (**Figures 1B-1C**). Particularly, genes associated with definitive hematopoiesis were up-regulated in Evi1^+^ fraction relative to the Evi1^−^ fraction; while genes related to differentiation, including T- and B-cell receptor signaling and T-cell differentiation, were upregulated in the Evi1^−^ fraction (**Figures 1D-1E**). These results suggested that the Evi1^+^ fraction might be enriched in cells associated with maintenance of hematopoiesis, while the Evi1^−^ fraction might contain cells associated with the early stages of lineage commitment.

To investigate this possibility further, we also sought to compare gene expression patterns in the Evi1^+^ and Evi1^−^ fractions at the single cell level. We profiled a panel of 96 genes associated with HSC-self-renewal and lineage commitment using a real-time single cell qPCR array (96*·*96 Dynamic Array, BioMark™ system, Fluidigm). A detailed gene list and primers used are provided in the **Supplemental Table S1**.

Consistent with our microarray data we observed down-regulation of negative markers of HSC quiescence such as *Cd48* and *Cdk6* (Kiel et al. 2005, Scheicher et al. 2015), and up-regulation of quiescent HSC markers such as *Epcr, Ndn* and *Foxo1* (Balazs et al. 2006, Kubota et al. 2009, Tothova et al. 2007) in the Evi1^+^ fraction (see **Supplemental Figure S1A**), suggesting that the Evi1^+^ population is enriched in quiescent LT-HSCs.

We also observed distinct and significant differences between the two populations in the variability of gene expression. Using a multivariate analogue of Levene’s test for equality of variances – which uses the dispersion (see **STAR Methods**) as a multivariate measure of cell-cell variability that takes into account the variability of each gene as well as the patterns of co-variance between genes (Van Valen 2005) – we observed that the Evi1^−^ fraction is significantly more heterogeneous in its overall gene expression patterns than the Evi1^+^ fraction (**Figures 1F-1G**).

Taken together, our bulk and single cell profiling experiments suggest that Evi1^+^ fraction represents a more highly purified population than the Evi1^−^ that more consistently expresses markers of definitive hematopoiesis, and is enriched for LT-HSCs.

To test this possibility further, we sought to assess the *in vivo* function of the Evi1^+^ and Evi1-GFP^−^ fractions. Competitive bone marrow transplantation (BMT) indicated that functional LT-HSCs were indeed enriched in the Evi1^+^ population (**Figure 1H**). To quantify this enrichment we performed limiting dilution BMT. This analysis estimated that the expected incidence of functional LT-HSCs to be approximately 1 in every 6.7 cells in the Evi1^+^ fraction, compared to 1 in every 514.6 cells in the Evi1^−^ fraction, representing a 76.7-fold enrichment upon positive selection for Evi1 expression (**Figure 1I**). Collectively, these data indicate that Evi1^+^ fraction constitutes a purified population of functional LT-HSCs.

### The niche factor Angiopoietin-1 has widespread effects on HSC gene expression patterns

Once we had identified a homogeneous LT-HSC population, we sought to investigate the mechanisms of HSC self-renewal using a bespoke paired daughter cell (PDC) assay. Classical PDC experiments seed individual hematopoietic stem/progenitor cells into isolated wells *in vitro* and allow them to divide. Upon division, daughter cells are separated with a micro-manipulator and the colony forming ability of the daughter cells are compared by assessing the sizes and multi-lineage composition of the resulting colonies (usually after 7 or 14 days) (Takano et al. 2004, Ema et al. 2000). Typically, colony sizes are observed to follow an exponential distribution, with lineages apparently randomly distributed among colonies (Nakahata et al. 1982, Suda et al. 1984). Exponential clone size distributions are predicted by stochastic models of proliferation (Greulich & Simons 2016) and these two features are taken as evidence of inherent stochasticity in the original division that gave rise to the two daughter cells. However, it is possible that the observed stochasticity is due to culture conditions that do not accurately mimic the *in vivo* stem cell niche, or due to the complex spatio-temporal patterns of cell-cell interactions that arise during colony development, rather than from stochasticity in the initial stem cell division *per se*. To resolve this issue, we designed a PDC assay that compares daughter cell identities immediately after HSC divisions, and used this assay to explore the effects that niche conditions and cellular aging have on division patterns.

A schematic of our PDC assay procedure is given in **Figure 2A**. Briefly, individual Evi1^+^ LT-HSCs were seeded on to fibronectin (FN)-coated plates and cultured in serum-free medium with Stem Cell Factor (SCF) and Thrombopoietin (TPO), with or without the niche factor Angiopoietin-1 (Angpt1), which has previously been shown to enhance HSC quiescence and self-renewal (Arai et al. 2004), for two days to allow them to divide. After two days of culture, wells that contained precisely two cells were identified, daughter cells were separated using a micro-manipulator, and immediately profiled for expression of the same panel of 96 HSC-self-renewal and lineage commitment associated genes we used to compare Evi1^+^ and Evi1^−^ populations (96*·*96 Dynamic Array, BioMark™ system, Fluidigm, see **Supplemental Table S1** for a detailed list of genes and primers used). Because gene expression patterns for both daughter cells from each single LT-HSC division were obtained, we were able to compare their patterns of gene expression, and explore how multivariate associations between daughter cells were affected by culture conditions.

**Figure 2.**
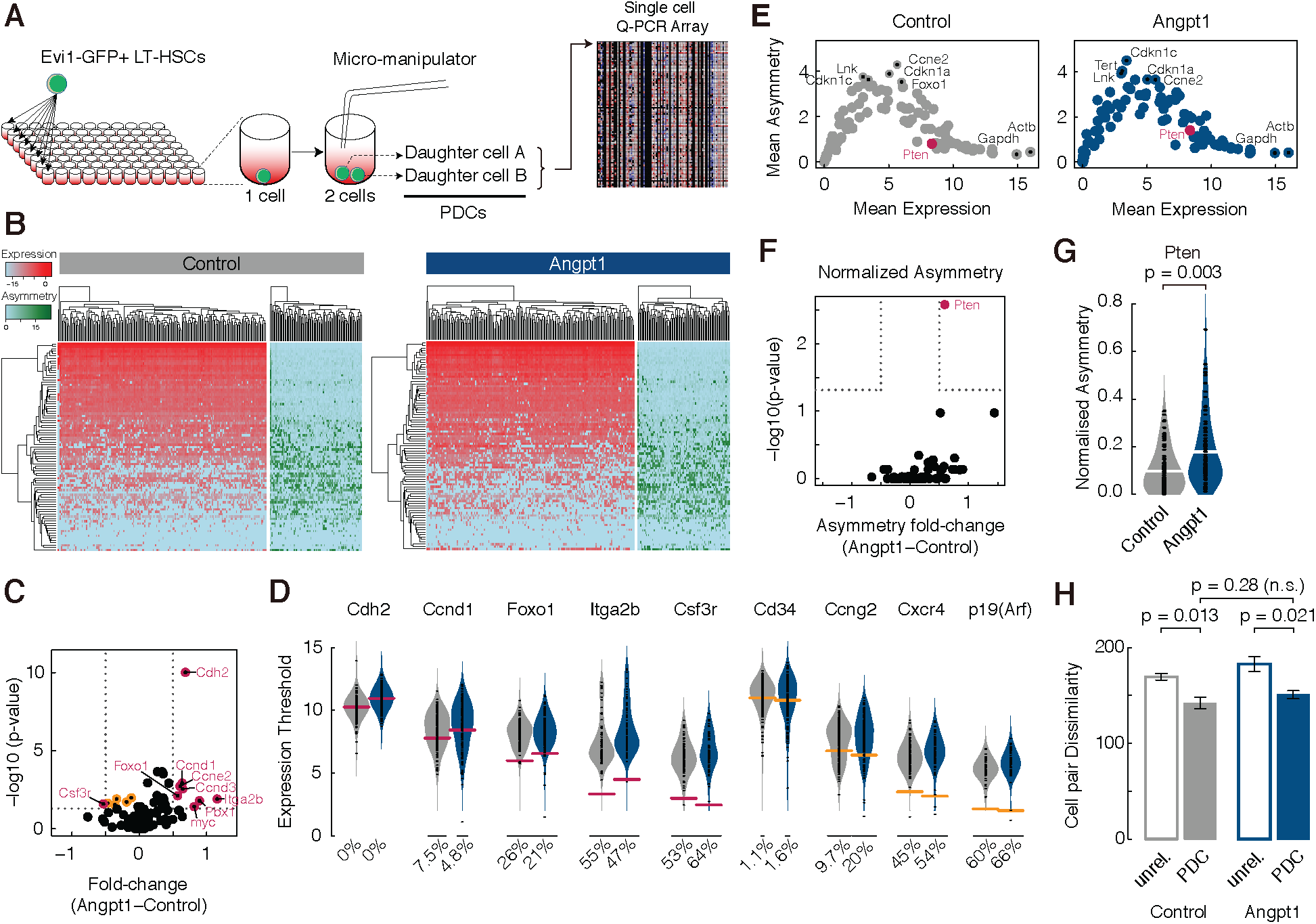
Combinatorial effects of Angpt1 treatment on cell division asymmetry. (A) Paired-daughter cell assay. Individual Evi1^+^ cells were seeded on to multi-well plates. Following cell division *in vitro*, daughter cells were manually separated and processed for gene expression analysis by qPCR array. (B) Heat-map of single-cell gene expression and asymmetry patterns from control (left) and Angpt1 treated (right) cultures. Expression patterns are in red-blue, asymmetry patterns are in green-blue. (C) Volcano-plot, showing the expression difference between treatment conditions (log-fold-change of mean) and negative log of p-values (FDR corrected using the Benjamni-Hochberg [BH] method) from a hypothesis test for single cell differential gene expression (McDavid et al. 2013). Significant genes (FDR corrected p-value < 0.05, and expected fold change > 1.4) are highlighted in cerise; orange points mark significant genes that do not meet the fold-change criterion. (D) Violin-plots of expression values in control (grey) and treatment (blue) conditions for selected genes. Percentage values along the x-axis indicate the proportion of zero readings. Note that the proportion of zeros can vary substantially and thereby contribute considerably to statistical significance. Cerise and orange bars indicate expression mean, and fold-change levels from panel C. (E) Plots of mean expression against mean division asymmetry in control (left, grey) and Angpt1 treated (right, blue) LT-HSCs. (F) Volcano-plot, showing the asymmetry difference between treatment conditions (log-fold-change of mean asymmetry) and negative log of p-values (FDR corrected using the BH method). Significant genes (FDR corrected p-value < 0.05, and fold change > 1.4) are highlighted in cerise. (G) Violin-plot of expression asymmetry for *Pten*. p-value is from a Wilcoxon rank sum test (FDR corrected using BH method). White bar indicates median. (H) Dissimilarity (assessed using L1-norm) of randomly selected daughter cell pairs (i.e. unrelated) and paired-daughter cells from the same division. Data is presented as mean ± standard deviation, p-values are from two sided t-tests (from three biological replicates containing 48 pairs of cells each).

To begin, however, we assessed patterns of gene expression without regard to pairing between daughter cells. Unsupervised clustering of this data indicated that patterns of gene expression were highly stochastic in both control and Angpt1 treated cells (**Figure 2B**). We observed that Angpt1 treatment significantly increased expression of cell adhesion molecules (such as *Cdh2, Itga2b*) responsible for HSC homing, and significantly reduced the expression of mobilization signal-receptors (such as *Csf3r, Cxcr4*) (Petit et al. 2002) and mobilization indicators (such as *Cd34*) (Tajima et al. 2000) (**Figures 2C-2D**). In parallel, we also observed increased expression of cell cycle regulators (such as *Ccnd1, Ccnd3, Ccne1, Foxo1, Ccng2*), and reduced expression of senescence factors such as *p19-Arf* following Angpt1 treatment (**Figures 2C-2D**). These results are in agreement with previous findings that Angpt1 improves cell survival (Arai et al. 2004), and indicate that Angpt1 regulation of HSC dynamics is highly combinatorial, affecting multiple regulatory pathways.

### Angpt1 has combinatorial effects on gene expression asymmetry

To investigate the effect of Angpt1 on division patterns we then compared gene expression profiles of paired daughters. Consistent with the stochastic nature of gene expression, we observed that expression asymmetry (defined as the difference in the expression of a given gene between the two daughters, see **STAR Methods** for more details) was also highly stochastic (**Figure 2B**). Notably, we observed that expression asymmetry had a nonlinear relationship to expression: those genes that had a high or low average expression tended to be symmetrically partitioned, while those that were sporadically-expressed were substantially more asymmetrically partitioned (**Figure 2E**). The most asymmetrically partitioned genes in both control and Angpt1 treated conditions included cell cycle regulators such as *Cdkn1c, Cdkn1a, Ccne2* and regulators of self-renewal such as *Lnk* (Ema et al. 2005). Despite the combinatorial effects of Angpt1 on gene expression patterns we found that only one gene – *Pten* a negative regulator of the PI3K-AKT signalling pathway that controls HSC proliferation, survival, differentiation, and migration (Yilmaz et al. 2006) – was deferentially partitioned on Angpt1 treatment (**Figures 2F-2G**).

To better understand the collective effect of expression changes on divisions we also determined the total asymmetry of each division (defined as the sum of the asymmetries for all genes profiled). Consistent with the per-gene analysis, we found that Angpt1 did not have a significant effect on overall division asymmetry (**Figure 2H**). However, in accordance with their common parental origin, we did observe that daughter cells from the same division tended to be more like each other than randomly selected pairs of unrelated cells from the daughter cell population. This daughter cell similarity was consistently observed in both control and Angpt1 treated conditions (**Figure 2H**).

Collectively, these results indicate that Angpt1 has a widespread, combinatorial affect on both gene expression and expression asymmetry patterns. However, while Angpt1 did not appear to affect overall division asymmetry we were not able to assess if it altered the balance between symmetric self-renewal divisions (which produce two stem cells) and symmetric differentiation divisions (which produce two committed, or partially committed, daughter cells) using unsupervised statistical analyses. Yet, regulation of this balance is critical to stem cell homeostasis (for a mathematical argument of why this balance is crucial, see **Supplemental Material: Mathematical Modeling**). To investigate further we therefore sought to develop a supervised machine learning approach to extract more detailed information about the identities of the daughter cells produced from our PDC experiments.

### Machine learning of cell identities from single cell expression data

Our unsupervised analysis suggested that single cell gene expression patterns in LT-HSCs are highly stochastic (**Figure 2B**). We reasoned that some of this variability was biologically meaningful yet, some arises due to inherent technical limitations of single cell profiling methods, which confound statistical analysis. To better understand the nature of this variability we therefore developed a supervised machine learning approach to discriminate biological from technical noise.

To do so, we first constructed a training library of single cell expression patterns from which we were able to benchmark the expected cell-cell variability in expression patterns that arises when profiling various different defined cell populations. Specifically, we isolated a range of different types of hematopoietic stem and progenitor cells from young (4-week old), adult (8-week old) and aged (18-month old) mice. In total we selected nine different stem and progenitor populations. Details of our selection procedures are given in **Figure 3A** and the **STAR Methods**. Details of the abbreviations used to identify these populations are given in the **STAR Methods**. Immediately after isolation we profiled these populations for the same panel of 96 HSC-related genes that we used in the PDC experiments using the same single cell qPCR array (96*·*96 Dynamic Array, BioMark™ system, Fluidigm, see **Supplemental Table S1** for a full list of genes and primers used). Thus, we obtained samples of single cell expression patterns for a range of cell types in the hematopoietic hierarchy, taken from mice at different stages of development. These sample data sets provided us with a training library from which we were able to learn the natural variation in gene expression present in each of these populations, and how gene expression patterns compare across the populations and ages.

**Figure 3.**
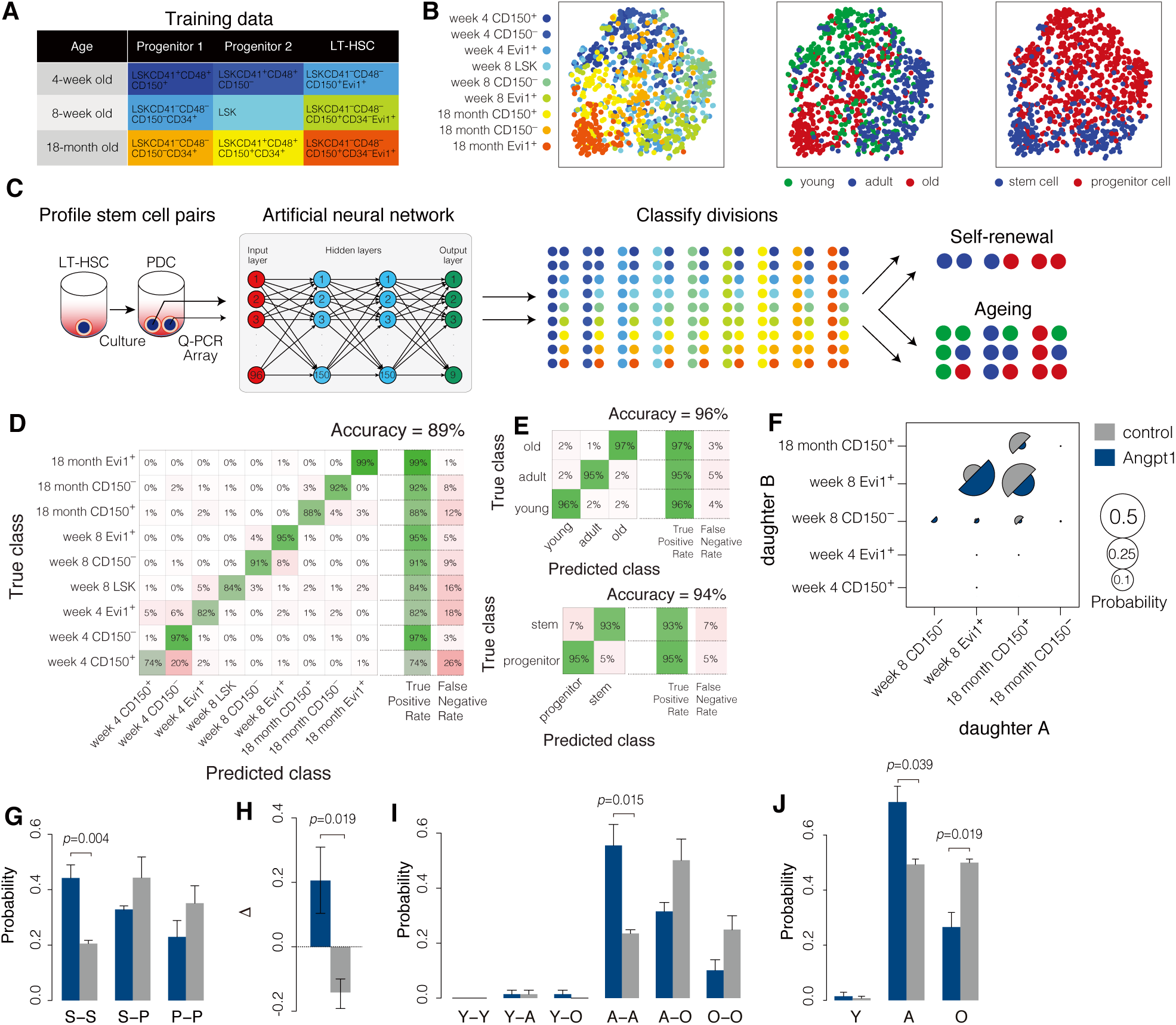
Machine learning of cell identities from single cell expression data. (A) Selection protocols for training data. (B) Dimensionality reduction of training data using t-SNE (Van Der Maaten et al. 2009). Training data sets exhibit broad patterns of gene expression and are not easily distinguished from one-another with unsupervised methods. (C) Schematic of machine learning protocol. An artificial neural network (ANN) was trained to distinguish between cells from training samples based on their patterns of gene expression. Once trained the ANN was then used to predict the identity of paired daughter cells (PDCs). (D-E) The trained ANN is able to reliably identify cells from their expression patterns. (D) Confusion matrix for the ANN classifier, indicating high (89%) accuracy. (E) When restricted to age (confusion matrix, top) or regenerative status (confusion matrix, bottom) classification accuracy further improves. (F) Classification of PDC divisions using trained ANN. Daughter cell identities are not ordered. (G) Classification of PDC divisions using the trained ANN, restricted to regenerative status. (H) The expansion factor Δ. When Δ > 0 the stem cell pool is expanding; when Δ < 0 the stem cell pool is depleting. (I-J) Angpt1 treatment inhibits cellular aging *in vitro*. (I) Classification of PDC divisions using trained ANN, restricted to age. Spontaneous aging is commonly observed in one or other, or both, daughter cells. Spontaneous rejuvenation is never observed, to within accuracy of classifier. (J) Probability distribution of daughter cell ages without regard for pairings. In all panels data is presented as mean ± standard deviation (from three biological replicates of 48 pairs of cells in each condition); p-values are from two-sided t-tests.

Unsupervised analysis of this training data indicated that each population was highly variable in its expression patterns (**Figure 3B**). While distinct trends were apparent – for example cells taken from mice of the same age tended to be similar to each other, see **Figure 3B, middle panel**; as were stem cells from mice of all ages, see **Figure 3B, right panel** – these associations were diffuse and highly combinatorial, making it difficult to discriminate between the populations using traditional statistical methods.

To progress further, we trained an artificial neural network (ANN) to accurately discriminate between the different training cell types based on their patterns of expression. ANNs are highly effective at pattern recognition and can significantly out-perform classical statistical procedures when properly trained (Friedman et al. 2001, Goodfellow et al. 2016). Details of the architecture and training of our ANN are given in the **STAR Methods** and summarized schematically in **Figure 3C**.

We found that our ANN was able to recognize all the different kinds of stem and progenitor cells profiled, and simultaneously predict the age of the mouse from which they were isolated, with 89% overall accuracy. The confusion matrix for cross-validated predictions is given in **Figure 3D**. When restricted to predicting the regenerative status of the cell (i.e. is it a stem or progenitor cell?) or predicting the age of the cell (i.e. was it taken from a young, adult or aged mouse?), accuracy increased to 96% and 94% respectively. Confusion matrices for cross-validated predictions of regenerative status and age are given in **Figure 3D** and **Figure 3E**.

These results show that although gene expression patterns are highly stochastic, and variations in expression patterns between different types of stem and progenitor cells are highly combinatorial, machine learning methods can be used to benchmark this variability and accurately predict individual cell identities directly from their gene expression profiles.

### Hematopoietic stem cell differentiation is niche-regulated yet inherently stochastic

To investigate the nature of HSC divisions further we then used our trained ANN to determine the identities of the daughter cells produced from our PDC experiments. Because our ANN assigns each daughter cell to one of nine different categories, we were able to classify each division into one of 81 (9 × 9) different categories, representing a range of different kinds of symmetric and asymmetric divisions, as well as predict the developmental ‘age’ of each daughter cell to determine if any aging events had occurred through division (**Figure 3C**).

By analyzing PDC division patterns we found that LT-HSCs taken from 8-week old mice typically divided to produce a range of daughter cells with different regenerative potentials and predicted ages (**Figure 3F**). In both control and Angpt1 treated conditions the most common divisions produced were ones in which: (1) both daughter cells identified as belonging to the Evi1^+^ population from 8-week old mice. These divisions represent symmetric stem-stem (S-S) divisions in which both daughter cells retain the same regenerative status and age characteristics of the parent LT-HSC; (2) one daughter cell is classified as an 8-week old Evi1^+^ LT-HSC, and one daughter is classified as an 18-month old LSKCD41^+^CD48^+^CD34^+^CD150^+^ progenitor cell. These divisions represent asymmetric stem-progenitor (S-P) divisions in which one daughter cell retains the same regenerative status and age characteristics of the parent LT-HSC and one daughter acquires a partially differentiated status and characteristics associated with aging; (3) both daughter cells are classified as 18-month old LSKCD41^+^CD48^+^CD34^+^CD150^+^ progenitor cells. These divisions represent symmetric progenitor-progenitor (P-P) divisions in which both daughters differentiate and both acquire characteristics associated with aging.

However, while similar kinds of division were seen in both culture conditions, the balance of probabilities of these divisions occurring was significantly altered upon Angpt1 treatment (**Figures 3G-3J**). Considering the regenerative status of the daughter cells (without regard for their predicted developmental age) we found that the total proportion of cells undergoing asymmetric S-P self-renewal divisions was not affected by Angpt1 treatment. This is in accordance with our previous unsupervised analysis of gene expression patterns, which found that Angpt1 did not affect the symmetry of divisions (**Figure 2**). However, our ANN analysis revealed that the proportion of symmetric S-S divisions was significantly increased by Angpt1 treatment (**Figure 3G**), while the proportion of symmetric P-P divisions was decreased upon Angpt1 treatment, a feature that was not apparent using unsupervised methods. These results suggest that while Angpt1 treatment does not affect overall propensity for symmetric divisions, it might affect stem cell numbers by altering the balance between different kinds of symmetric division.

This is notable because a simple mathematical analysis of cell proliferation dynamics (see **Supplemental Information: Mathematical Modeling**) indicates that maintenance of the stem cell pool is critically determined by the relative propensity of symmetric S-S and P-P divisions. In particular, if the probability *p*_1_ of an S-S division is greater than the probability *p*_2_ of a P-P division then the stem cell pool will expand. Similarly, if *p*_2_ is greater than *p*_1_ then the pool will deplete. Thus, the rate of expansion or depletion of the stem cell pool is determined by the difference Δ = *p*_1_ - *p*_2_. We found that the expansion parameter Δ < 0 in control conditions, yet Δ > 0 upon Angpt1 treatment (**Figure 3H**), indicating that Angpt1 treatment may sustain stem cell self-renewal *ex vivo* by specifically stimulating symmetric S-S divisions.

We also used our ANN to determine composition of the Evi1^−^ population. Consistent with our previous analysis of the relative variability of the Evi1^+^ and Evi1^−^ populations, our ANN classifier predicted that the Evi1^−^ fraction contains a mixture of stem and progenitor cell populations (**Supplemental Figures S1B-S1C**).

### *Ex vivo* culture induces spontaneous cellular aging from the first division

In addition to predicting regenerative status our ANN was also able to predict the developmental ‘age’ of cell. To investigate how culture affects cellular aging we therefore also used our ANN to investigate the predicted ages of paired daughter cells. Depending on its similarity to cells in the training library we classified each daughter cell as being young (most similar to a cell taken from a 4-week old mouse, denoted Y), adult (most similar to a cell taken from an 8-week old mouse, denoted A) or old (most similar to a cell taken from an 18-month old mouse, denoted O), and thus classified each LT-HSC division into one of nine possible classes depending on the identities of the two daughter cells: young-young (Y-Y), young-adult (Y-A), young-old (Y-O), adult-adult (A-A), adult-old (A-O), or old-old (O-O).

We first noted that while it is theoretically possible for daughter cells from divisions of LT-HSCs taken from 8-week old adult old mice to acquire a younger developmental age in culture, such spontaneous rejuvenation did not occur in practice (**Figure 3I**). However, while both daughters were commonly identified as being adult cells, we also observed that spontaneous aging of one or other (or both) daughter cells also occurred with significant regularity (**Figure 3I**). This indicates that cell culture induces accelerated cellular aging in the progeny of LT-HSCs from the very first division. Indeed, in control conditions approximately 50% of cells showed spontaneous aging through the first division (**Figure 3J**). However, we also observed that this spontaneous aging was significantly inhibited upon Angpt1 treatment (**Figures 3I-3J**), suggesting that it is a feature of culture that is not inevitable, but can be controlled. This analysis highlights the need for culture conditions that are capable of regulating both regenerative status and cellular aging.

### HSC self-renewal ability and sensitivity to niche instruction is rapidly lost in culture

Next, we wanted to determine if the effect of Angpt1 on division patterns that we observed was passed though multiple cell divisions. To do so we examined associations between paired granddaughter cells (PGDCs). In this assay we isolated Evi1^+^ LT-HSCs from 8-week old mice, seeded them one-per-well onto FN-coated plates and cultured them in serum-free medium with SCF and TPO, with or without Angpt1 for two days. Wells that contained precisely two cells were identified and daughter cells from these first divisions were separated with a micro-manipulator, re-plated cultured for two more days and allowed to divide a second time in the same culture conditions. As before, wells that contained precisely two cells were identified, and granddaughters were separated and profiled for expression of the same panel of 96 HSC-self-renewal and lineage commitment associated genes we used previously (96*·*96 Dynamic Array, BioMark™ system, Fluidigm, see **Supplemental Figure S2A**).

Consistent with the results of the PDC assay we observed that almost all secondary divisions produced granddaughter cells that had lost stem cell status (**Supplemental Figure S2B**), and more than 70% of secondary divisions produced at least one granddaughter cell with aged characteristics in both control and Angpt1 treated cultures (**Supplemental Figure S2C**). However, in contrast to the PDC results, we found that Angpt1 did not significantly alter secondary division patterns, either with respect to age or regenerative status. Notably, because the propensity for P-P divisions dominated the propensity for S-S divisions in secondary divisions, we found that the expansion parameter Δ = *p*_1_ − *p*_2_ was negative for secondary divisions in both control and Angpt1 treated conditions, indicating that neither condition is able to support sustained stem cell self-renewal beyond one division.

Collectively, these results suggest that freshly isolated HSCs retain an intrinsic sensitivity to external stimuli which allows them to respond to supplementation of niche factors *in vitro*. However, they rapidly lose this sensitivity, and simultaneously gain aged characteristics, when cultured *ex vivo*.

### Substrate stiffness regulates LT-HSC division patterns

To investigate HSC division dynamics further we also sought to determine how the physical structure of the niche affects HSC behaviour. To do so we manufactured a bespoke polyethylene glycol (PEG) microwell culture system, which provides a softer environment with a high water content that more closely matches the physical characteristics of the *in vivo* HSC niche than standard culture conditions (Gobaa et al. 2011, Lutolf et al. 2009, Roch et al. 2017). Similar niche constructs have been shown to support stem cell self-renewal in other contexts (Gilbert et al. 2010).

To test the effect of our artificial niche on division patterns we conducted our PDC assay using 8-week old Evi1^+^ LT-HSCs seeded onto our FN-coated PEG hydrogel micro-wells with or without Angpt1. In the absence of Angpt1 we found that this culture system stimulated symmetric S-S divisions, resulting in a net expansion of the HSC pool (**Supplemental Figures S3A-S3C**), indicating that it is more supportive of HSC self-renewal than standard culture conditions. Notably, Angpt1 treatment did not significantly alter division patterns, perhaps because symmetric S-S divisions were already enhanced by the PEG micro-well niche. In accordance with our results from standard culture conditions we observed spontaneous aging, albeit at a lower rate, with approximately 20-30% of cells adopting an aged phenotype after one division in micro-well culture (**Supplemental Figures S3D-S3E**). However, in contrast to standard culture conditions, we also observed spontaneous rejuvenation of a small proportion of cells in both control and Angpt1 treated cultures, and saw that Angpt1 treatment appeared to weakly inhibit the aging process, significantly reducing the proportion of divisions that gave rise to two daughter cells with aged characteristics (**Supplemental Figure S3D**).

These results indicate that freshly isolated LT-HSCs are sensitive to both physical and biochemical stimulation by the niche, and variation of the physical characteristics of the culture environment may overcome the culture-induced loss of sensitivity to niche instruction and premature aging that we observed in standard culture conditions.

### Angpt1 increases LT-HSC numbers in vivo

Taken together our results suggest that Angpt1 treatment *in vitro* partially protects LT-HSCs from cellular aging, and increases the propensity for symmetric S-S divisions that expand the HSC pool. To investigate this further we sought to assess the regenerative capacity of paired daughter cells *in vivo*. To do so we performed limiting dilution bone marrow transplantation (BMT) of PDCs. As before, individual Evi1^+^ LT-HSCs were seeded onto FN-coated plates and cultured in serum-free medium with SCF and TPO, with or without Angpt1 for two days. After two days of culture, daughter cells were collected and 1, 2, 3, 4, or 6 sets of PDCs were transplanted into lethally irradiated mice, and the proportion of LT-HSCs in the transplanted daughter cells was estimated from the fraction of non-responding mice (**Figure 4A**). Consistent with the results of our ANN classifier, we observed that culture with Angpt1 increased the proportion of LT-HSCs nearly 6-fold, from 1 in 58.6 transplanted cells without Angpt1 treatment to 1 in 9.7 with (**Figure 4B**). A similar analysis using PDCs cultured in our bespoke PEG micro-well also confirmed the enhanced ability of this culture system to maintain stem cell numbers *ex vivo* (**Supplemental Figure S4**).

**Figure 4.**
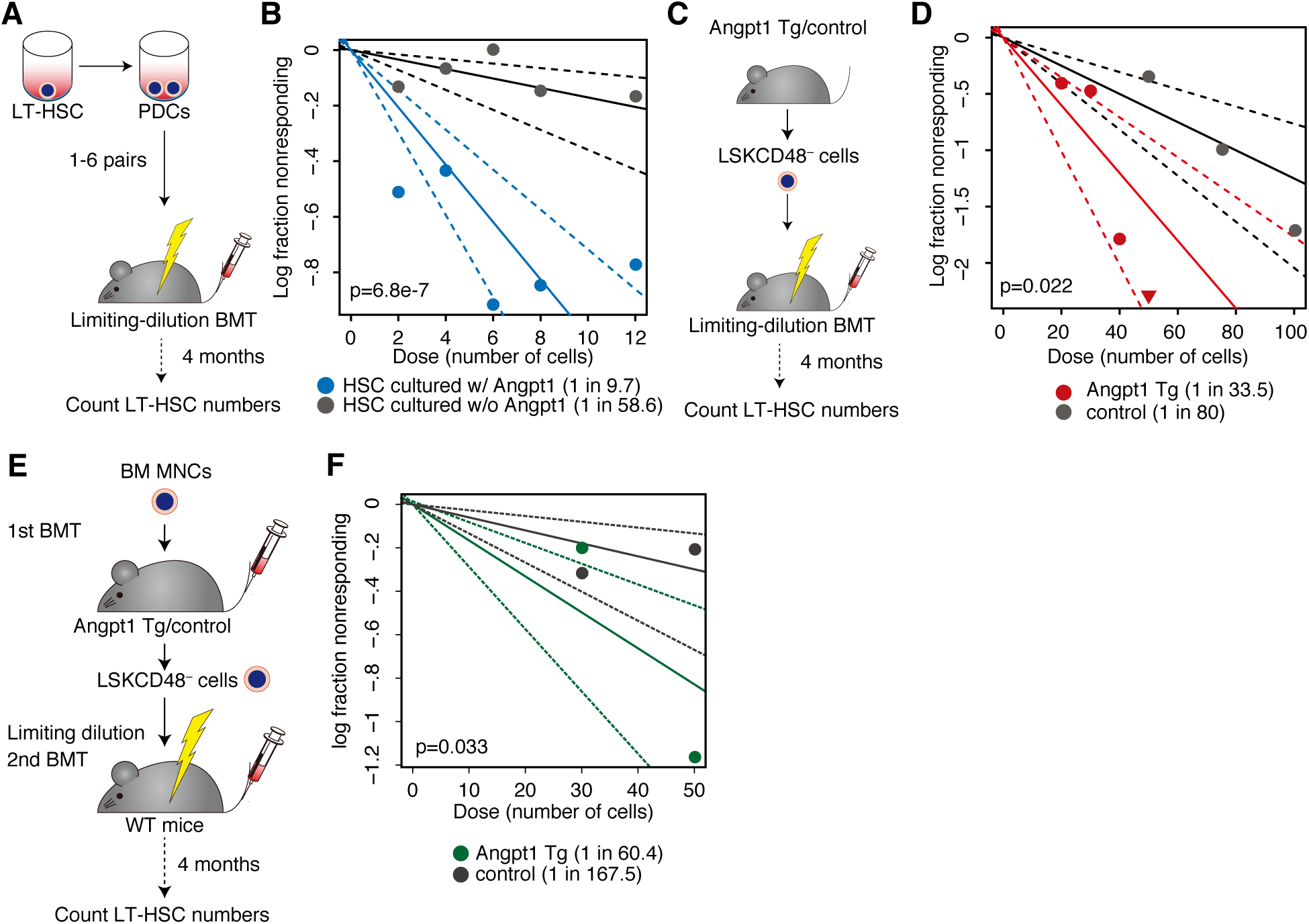
Angpt1 enhances functional stem cell activity *ex vivo* and *in vivo*. (A) Schematic of limiting dilution bone marrow transplantation (BMT) assay. (B) Limiting dilution BMT of cultured PDCs shows that Angpt1 treatment maintains HSC activity *ex vivo*. (C) Schematic of limiting dilution BMT assay in osteoblast-specific Angpt1 transgenic (Tg) mice. (D) Limiting dilution BMT in osteoblast-specific Angpt1 Tg mice shows that Angpt1 enhances HSC activity *in vivo*. (E) Schematic of secondary transplantation assay in osteoblast-specific Angpt1 Tg mice. (F) Secondary BMT of LSKCD48^−^ cells from osteoblast-specific Angpt1 Tg mice shows that Angpt1 increases HSC numbers *in vivo*. p-values are from extreme value limiting dilution analysis (ELDA) (Hu & Smyth 2009) in all panels.

In addition to determining the effect of Angpt1 on cultured cells we also evaluated its role in regulating LT-HSC numbers *in vivo*. To do so, we isolated LSKCD48^−^ cells from osteoblast-specific Angpt1 transgenic (Tg) (Col1a1-Cre(+); CAGp-IND-COMP-Angpt1) and control (Col1a1-Cre(–); CAGp-IND-COMP-Angpt1) mice and assessed the regenerative function of these cells via limiting dilution BMT (**Figure 4C**). By analyzing the fraction of non-responding mice subsequent to BMT we were able to estimate the proportion of LT-HSCs in the LSKCD48^−^ fraction. We found that LT-HSC incidence was significantly higher in Angpt1 Tg mice than controls (1 in 33.5 cells for Tg mice, compared to 1 in 80 for controls, see **Figures 4C-4D**).

To further validate these results we also performed secondary transplantation. We isolated and transplanted BM mononuclear cells into osteoblast-specific Angpt1 Tg and control mice. Four months after primary BMT we isolated donor-derived LSKCD48^−^ cells, performed further limiting dilution BMT, and again estimated LT-HSC numbers from the fraction of non-responding mice. We found that donor-derived cells isolated from Angpt1 Tg mice were significantly enriched for LT-HSCs in comparison to donor-derived cells isolated from control mice (1 in 60.4 cells for Tg mice, compared to 1 in 167.5 for controls, see **Figures 4E-4F**).

Immunohistochemical staining of VE-cadherin^+^ blood vessels in the BM indicated that total vessel length was comparable in Angpt1 Tg and control mice (**Supplemental Figure S5**), suggesting that Angpt1 does not boost stem cell numbers in Angpt1 Tg mice by enlarging the HSC niche. Rather, these results are in agreement with our PDC analysis and indicate that Angpt1 maintains stem cell activity *in vivo* by stimulating symmetric S-S divisions in the stem cell pool.

### Stem cell self-renewal potential declines with age

Collectively, our results indicate that divisions of HSCs in adult (8-week old) mice are inherently stochastic, yet regulated by both the biochemical and physical structure of the niche. We were interested to determine if this inherent stochasticity and niche sensitivity were characteristic of stem cells taken from mice of all ages, or alternatively, if stem cell dynamics evolve with age.

To do so, we conducted further PDC experiments using LT-HSCs from young (4-week old) and aged (18-month old) mice in standard culture conditions, and again used our ANN to investigate daughter cell characteristics.

#### Division patterns of young LT-HSCs

When we examined division pattern of LT-HSCs taken from 4-week old mice, we found that they differed significantly from those of HSCs taken from 8-week old mice. In both control and Angpt1 treated conditions the most common divisions of 4-week old HSCs were those that produced: (1) two daughter cells both identified as belonging to the Evi1^+^ population from 4-week old mice. These divisions represent symmetric S-S divisions in which both daughter cells retain the same regenerative status and age characteristics of the parent LT-HSC; (2) one daughter cell is identified as belonging to the Evi1^+^ population from 4-week old mice and the other is identified as belonging to the Evi1^+^ population from 8-week old mice. These divisions represent symmetric S-S divisions in which one daughter cell acquires characteristics associated with aging.

Considering the regenerative status of the daughter cells (without regard for their predicted developmental age) we found that symmetric S-S divisions predominated over S-P and P-P divisions (**Figure 5B**), resulting in a positive expansion parameter Δ (**Figure 5C**), in both control and Angpt1 treated conditions, indicating that young HSCs have an inherent ability to expand their numbers independent of the culture conditions. Perhaps because of this inherent bias toward S-S divisions, Angpt1 did not significantly affect division patterns (**Figure 5B**) or significantly affect expansion dynamics (**Figure 5C**).

**Figure 5.**
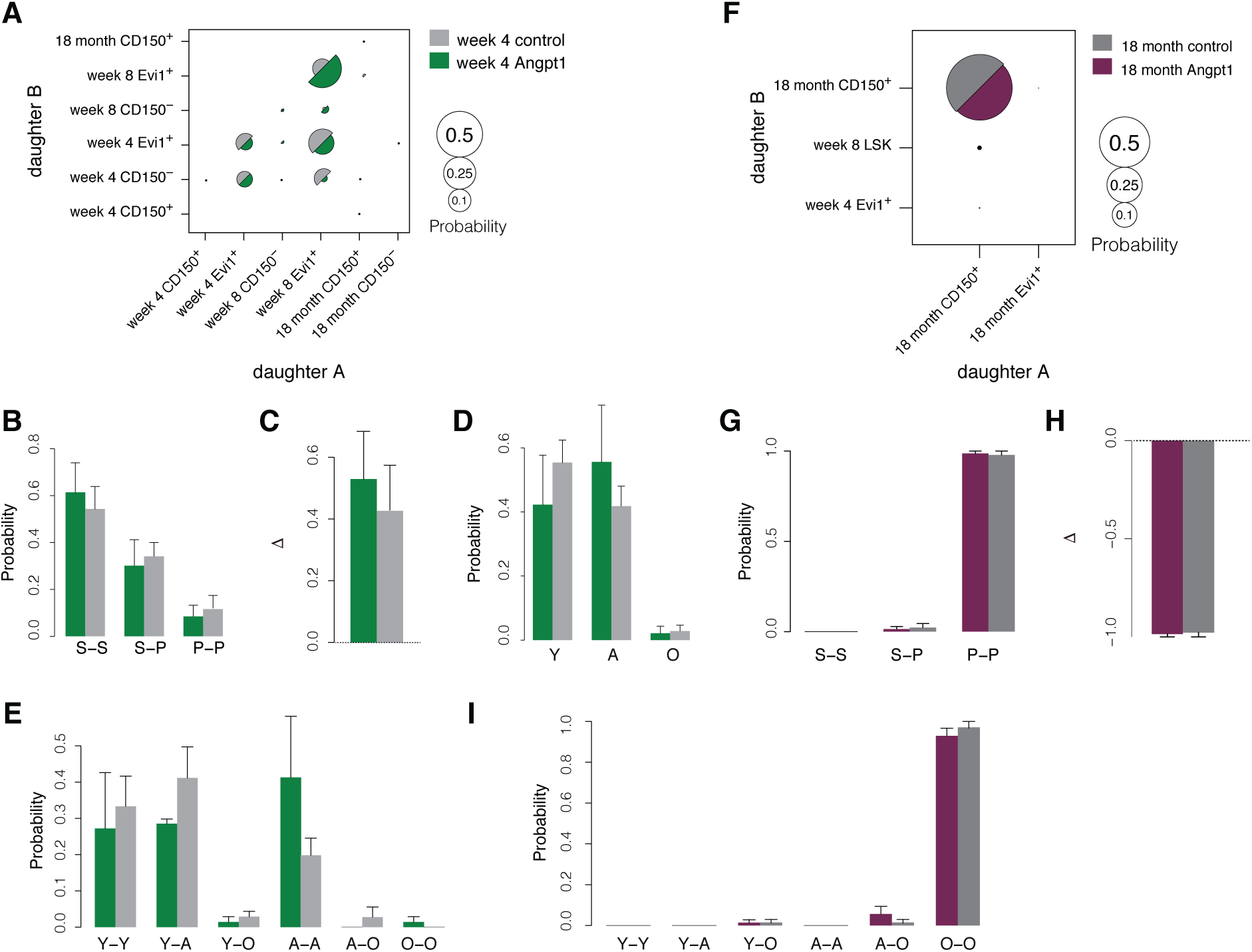
Machine learning classification of cell division patterns from young and aged stem cells. (A) Classification of divisions from young (4-week old) stem cells using trained ANN. Daughter cell identities are not ordered. (B) Classification of divisions from young stem cells using trained ANN, restricted to regenerative status. (C) The expansion factor Δ for young stem cells. Δ > 0 in both control and Angpt1 treated cells indicating robust expansion of the stem cell pool. (D) Probability distribution of predicted daughter cell ages using trained ANN from young HSC divisions without regard for pairings. (E) Classification of divisions from young stem cells using trained ANN, restricted to age. Spontaneous aging is commonly observed in one or other, or both, daughter cells in both control and Angpt1 treated conditions. (F) Classification of divisions from old (18-month old) stem cells using trained ANN. Daughter cell identities are not ordered. (G) Classification of divisions from old stem cells using trained ANN, restricted to regenerative status. (H) The expansion factor Δ for young stem cells. Δ < 0 in both control and Angpt1 treated cells indicating rapid depletion of the stem cell pool. (I) Classification of divisions from old stem cells using trained ANN, restricted to age. All cells, to within the accuracy of the classifier, have an aged phenotype in both control and Angpt1 treated conditions. In all panels data is presented as mean ± standard deviation (from three biological replicates of 96 cells in each condition for young cells and four biological replicates for old samples).

As with adult LT-HSCs we observed that young LT-HSCs underwent spontaneous aging from the first division in culture, with approximately 50% of divisions giving rise to a cell with adult (i.e. 8-week old) characteristics (**Figures 5A, 5D, 5E**). However, daughter cells did not readily take on characteristics of old (i.e. 18-month old) cells, suggesting that aging is a multi-step process that advances through multiple cell divisions. In accordance with its effect on regenerative status, Angpt1 treatment did not have a significant effect on the aging process in cells from young mice (**Figures 5D-5E**).

#### Division patterns of old LT-HSCs

In contrast to LT-HSCs taken from 4- and 8-week old mice, LT-HSCs taken from 18-month old mice divided almost exclusively to produce two cells with progenitor characteristics (**Figures 5F-5H**), resulting in a negative expansion parameter Δ (**Figure 5H**), suggesting that although these aged cells have the immunophenotypic characteristics of stem cells, they have lost the ability to self-renew *in vitro*. Furthermore, old LT-HSCs almost exclusively produced old daughter cells (**Figure 5I**), and we did not see spontaneous rejuvenation to a younger phenotype to within the error of the classifier. Very similar division patterns were observed in control and Angpt1 treated cultures, suggesting that, like young LT-HSCs, aged LT-HSCs are insensitive to niche instruction via Angpt1 *in vitro* (**Figures 5G-5I**).

In addition to using our ANN to investigate PDC patterns, we also used it to determine composition of the Evi1-GFP^−^ population from young and aged mice. We observed that the proportion of Evi1-GFP^−^ cells classified as Evi1^+^ increases with age, indicating that these two populations are initially quite distinct but become more alike in their gene expression patterns over time (**Supplemental Figure S1**).

Collectively, these results indicate that LT-HSC division patterns vary considerably with age, with regenerative potency being gradually lost, and response to Angpt1 being maximized through the middle of life. Notably, we found that the Angpt1 receptor Tie2 was consistently expressed in hematopoietic stem and progenitor cells throughout development and aging (**Supplemental Figure S6**), indicating that the observed changes in division patterns are not directly due to changes in sensitivity to niche instruction *per se* but rather might be due to an evolving balance between cell-intrinsic and extrinsic regulatory mechanisms.

### A mathematical model of HSC differentiation

To gain further clarity on this balance we sought to develop a mathematical framework to explain our experimental results. Because the precise molecular details underpinning stem cell aging are not known, we sought to develop a conceptual framework with which to understand aging and differentiation, rather than specifying a detailed molecular model. Our model is illustrated schematically in **Figure 6**. Brief mathematical details are given in **Box 1: Mathematical Model**, and a full description is provided in the **Supplemental Material: Mathematical Modeling**.

**Figure 6.**
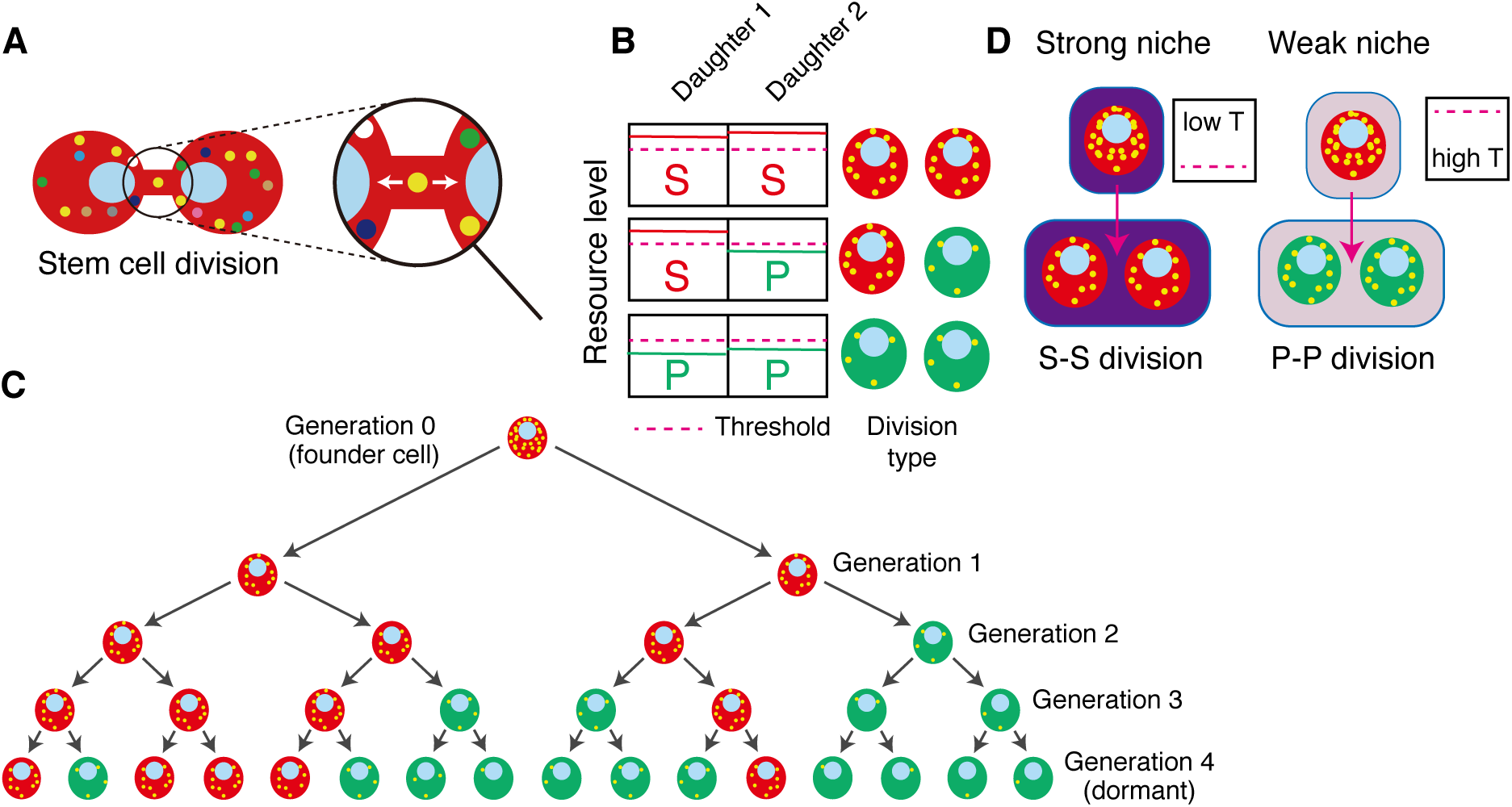
Schematic of mathematical model. (A) Each stem cell possesses a reservoir of a set of intracellular resources (here shown as differently colored dots) that are partitioned each time the cell divides. (B) Cell identity is determined by the abundance of intracellular resources relative to a threshold imposed by the niche. (C) If the resources are not fully replenished between divisions then the resource depletes with each division, and becomes increasingly likely to fall below the niche imposed threshold. (D) Different niches may impose different thresholds: when placed in a strongly instructive niche the threshold is set low, and the stem cell state can be maintained even if intracellular resources are low; when placed in a weakly instructive niche the threshold is set high, and the stem cell state can only be maintained if intracellular resources are high.

Our experimental results indicated that self-renewal ability gradually declines with age in a manner that is reminiscent of a linear decay or dilution process. To begin explain this loss, we first assumed that there is an intracellular resource (or set of resources) which define the stem cell identity. To account for niche instruction we then assumed that the identity of the cell is determined by the abundance of the intracellular resource relative to a threshold *T*_vivo_ imposed by the *in vivo* niche: if the resource abundance is above the threshold, then the cell adopts a stem cell identity; if the intracellular abundance is below the threshold then the cell differentiates. Thus, if the niche is strongly instructive (i.e. the threshold is *low*) then the stem cell identity can be maintained even if a cell’s internal resources are low; conversely if a cell has a high level of internal resources then it will maintain a stem cell identity even when niche instruction is weak (i.e. the threshold is *high*). Such threshold phenomena arise commonly in the regulatory molecular networks that define stem cell identities and are established, for example, by positive feedback mechanisms that stabilize alternate cell states (Alon 2006, MacArthur et al. 2009).

To account for aging we assumed that all adult HSCs arise from a common population of founder cells during development – for example, hemogenic endothelial precursor cells (Bertrand et al. 2010) – that possess an abundance of intracellular resource. However, each time one of these cells, or their progeny, divides the resource is partitioned stochastically (and possibly asymmetrically) between daughter cells. To account for intrinsic homeostasis we reasoned that the resource is likely replenished between divisions. If this replenishment occurs on a time-scale shorter than the cell cycle time, then the resource will be robustly maintained in the stem cell population, and differentiation will only rarely occur. However, if replenishment occurs on a time-scale longer than the cell cycle time, then the intracellular resource will become diluted with each successive division at a rate determined by the replenishment rate. In this case, it becomes increasingly likely that the resource abundance will fall below the threshold level as the stem cell population proliferates with aging. As it has previously been shown that HSCs have low protein turnover (Signer et al. 2014), we reasoned that slow replenishment dynamics may be characteristic of stem cells and the variation in division patterns with age that we observed might arise from such dilution phenomena.

### Ex vivo treatment with Angpt1 partially recapitulates the in vivo niche

The model described above accounts for stem cell dynamics *in vivo*. However, our experimental observations refer to division patterns *in vitro*. To model division patterns *in vitro* we assumed that culture could affect cell divisions by: (1) altering the symmetry of resource partitioning between daughter cells and (2) imposing a critical resource threshold that may be different to that encountered *in vivo* (i.e. *in vitro* culture conditions do not precisely mimic *in vivo* niche instruction).

We found that this simple conceptual model of stem cell aging explained the evolving division patterns we observed in both control and Angpt1 conditions with remarkable accuracy (**Figure 7A**). Culture conditions did not appear to significantly affect the dynamics of resource partitioning between daughter cells *in vitro* (our first assumption of *in vitro* dynamics, above). However, examination of model parameters indicated that Angpt1 treated conditions imposed a niche threshold that was substantially lower than control conditions yet higher than that encountered *in vivo*. This suggests that Angpt1 treatment improves culture conditions by producing an environment that partially recapitulates the *in vivo* niche.

**Figure 7.**
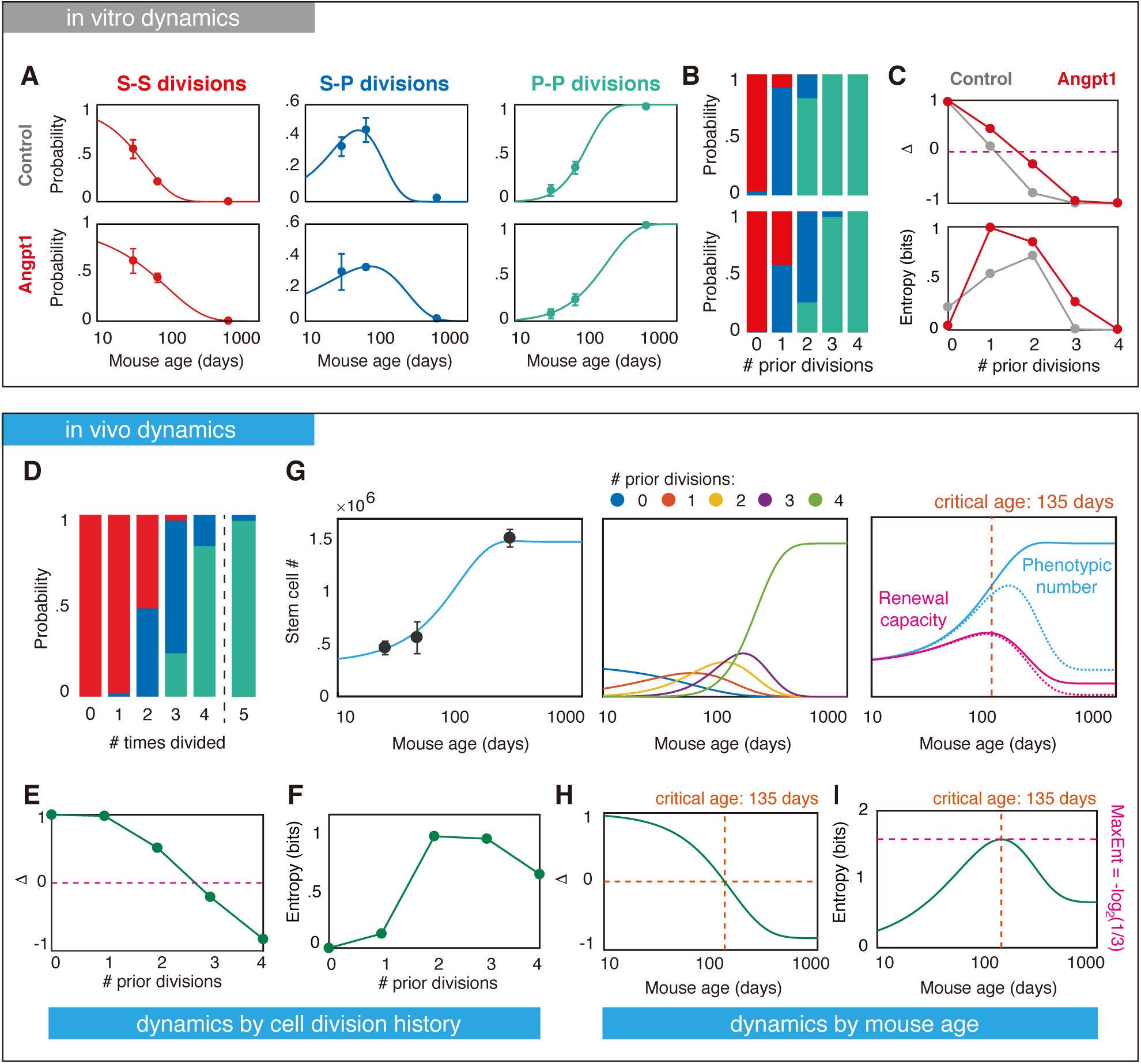
Cell division history determines HSC potency. (A) Fit of mathematical model to PDC division patterns. Markers show data; lines show model predictions. Division patterns in control cultures are shown on top, and division patterns in Angpt1 treated cultures are on the bottom. (B) Model estimates of the propensities for S-S, S-P and P-P divisions *in vitro* as a function of the number of times that a stem cell has divided previously (*in vivo*). Control (top), Angpt1 (bottom). (C) The division expansion factor Δ and entropy *in vitro* as a function of the number of times that a stem cell has divided previously. (D) Model estimates of the propensities for S-S, S-P and P-P divisions *in vivo* as a function of the number of times that a stem cell has divided previously. (E) The division expansion factor Δ *in vivo* as a function of the number of times that a stem cell has divided previously. (F) The division entropy *in vivo* as a function of the number of times that a stem cell has divided previously. (G) Left: plot of total stem cell numbers per mouse with age. Data are in black; model predictions, using parameters in panel D, are in blue. Middle: model estimate of total number of stem cells that have divided 0-4 times. Right: comparison of total stem cell number (blue) and renewal capacity (stem cell number multiplied by Δ, pink). Dotted lines show model predictions allowing for 5 stem cell divisions. (H) The division expansion factor Δ *in vivo* as a function of mouse age. At ∼ 135 days the stem cell population enters a phase of net decline. (I) The division entropy *in vivo* as a function of mouse age. At approximately 135 days, stem cell divisions are maximally stochastic. Pink line shows the entropy of the the maximum entropy uniform distribution over three states (for which the Shannon entropy is equal to log_2_(3)). In all panels data is presented as mean ± standard deviation (from at least three biological replicates).

Because our model incorporates cell cycle dynamics it allows translation from developmental time (i.e. mouse age) to cellular time (i.e. number of times divided, see **Supplemental Materials: Mathematical Modeling** for details of this translation). Therefore, model fitting to observed division patterns also allowed us to infer the propensity of an individual stem cell to undergo S-S, S-P and P-P divisions as a function of the number of times it had divided previously *in vivo* and the conditions it was cultured in (**Figure 7B**). We found that, by decreasing the niche threshold, Angpt1 treatment primarily acted to increase the propensity of stem cells that have divided one or two times previously to undergo S-S and S-P divisions respectively, yet has little effect on cells that have never divided or have divided three or more times (compare **Figure 7B**, top panel, to **Figure 7B**, bottom panel). The net consequence of this change is to increase the expansion parameter Δ, which quantifies self-renewal ability, for cells that have divided once or twice (**Figure 7C**, top panel), suggesting that Angpt1 specifically acts to enhance the self-renewal ability of this subset of the stem cell population.

##### Box 1: Mathematical model

Let *x*_*g*_ be a random variable denoting the copy number of the intracellular resource *X* in a stem cell that has divided *g* times previously and let *p*_*k*_(*g*) = *p*(*x*_*g*_ = *k*) denote the probability mass function of *x*_*g*_.

For ease of calculation we will consider a single founder cell in which the initial copy number of *X* is *n*. Thus, *p*_0_ = *p*(*x*_0_ = *k*) = *δ*_*kn*_, where *δ*_*i j*_ is the Kronecker delta function (*δ*_*i j*_ = 1 if *i* = *j* and zero otherwise). To account for niche regulation of cell divisions *in vivo* we assume that differentiation occurs if *x*_*g*_ falls below a critical threshold *T*_vivo_.

Upon cell division, the resource of the parent cell is passed onto the two daughters (denoted 1 and 2). For simplicity, we assume that *in vivo* each daughter inherits each copy of *X* with probability 1*/*2 (i.e. divisions are symmetric at the molecular level). To allow for resource replenishment we assume that subsequent to division each of the copy of *X* that is ‘lost’ to the sister cell is recovered with probability *q* before the next division. The parameter *q* then determines the rate of dilution of *X* in the population: if *q* = 0 then the resource in not replenished at all between divisions; while if *q* = 1 then the resource is fully replenished in each daughter between divisions. Under these assumptions the dynamics of *x*_*g*_ are governed by a Galton-Watson process (Van Kampen 1992) and *p*_*k*_(*g*) is a binomial distribution:

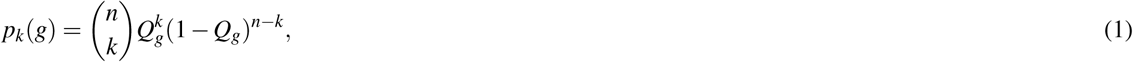

where 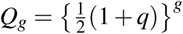 quantifies the extent to which the resource has been diluted after the *g*-th division.

Eq. (1) describes the dynamics of resource dilution in stem cells and their progeny *in vivo*. To investigate stem cell divisions *in vitro* we additionally assume that differentiation occurs if the resource level falls below a second critical threshold *T* that is determined by the culture condition and is different from that encountered *in vivo*. To allow for stress-induced changes to division patterns we also assume that daughter cell 1 inherits each copy of *X* with probability *α* and assume that the daughter cells are harvested before the resource has time to replenish. In the **Supplemental Material: Mathematical Modeling** we show that the probabilities *p*_11_(*g*), and *p*_00_(*g*) that a stem cell that has divided *g* times *in vivo* will perform an S-S or P-P division *in vitro* respectively are:

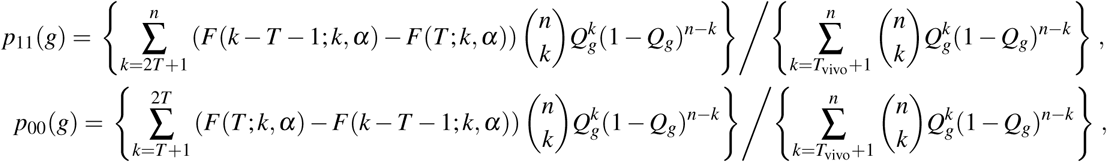

where *F*(*x*; *k, α*) is the cumulative distribution function for the binomial distribution B(*k, α*). The terms in the denominator of these equations ensure that the probabilities are only calculated over the stem cell pool (i.e. the probabilities are conditioned on the parent cell being a stem cell). The probability that a cell that has divided *g* times *in vivo* will perform an asymmetric division *in vitro* is *p*_01_(*g*) = 1 - *p*_00_(*g*) - *p*_11_(*g*).

Because these probabilities are not directly observed in experiment, we also require a model of the cell cycle to translate from cell intrinsic time (i.e. number of times divided previously) to developmental time (i.e. mouse age). Here, we assume that cell divisions within the stem cell pool are independent of one another and cell cycle times are Gamma distributed, a distribution that arises naturally from consideration of the cell cycle as a multi-stage Markov process, and has been observed experimentally (Yates et al. 2017). Full details of the translation between cell division number and developmental time are given in the **Supplemental Material: Mathematical modeling**.

### Angpt1 treatment increases division stochasticity

In addition to assessing how changes in the balance of divisions might affect cellular proliferation, we were also interested to identify if division stochasticity changed with age. To quantify division stochasticity we calculated the Shannon entropy of the probability distribution for S-S, S-P and P-P divisions as a function of number of times the parent cell had divided. Shannon entropy is a simple information-theoretic measure of the amount of uncertainty associated with an event: if the event is deterministic then there is no uncertainty in outcome, and the Shannon entropy is zero. Conversely, if many possible outcomes can occur with equal likelihood – as is the case when throwing a fair die for example – then the event is maximally stochastic and the Shannon entropy reaches a maximum of log_2_(*N*), where *N* is the number of events that can occur (Cover & Thomas 2012). Here, the Shannon entropy is an estimate of the uncertainty in the outcome of a division for a randomly selected stem cell from the pool.

We found that cells that had never divided almost always produced two stem cells on division in both control and Angpt1 treated conditions (**Figure 7B**). Thus, because there is low uncertainly in division patterns they have very low entropy (i.e. divisions are nearly deterministic). Similarly, those cells that had divided four times or more always produced two progenitor cells on division, in both control and Angpt1 treated conditions (**Figure 7B**). In this case, because there is no uncertainly in division patterns, they have zero entropy (i.e. divisions are perfectly deterministic). However, in contrast cells, that had divided 1-3 times previously were able to realize a range of division patterns, and so exhibited higher entropy, and therefore more stochasticity. Comparing the entropy of division patterns revealed that Angpt1 treatment typically increased division entropy, indicating that niche instruction via Angpt1 encourages more stochastic stem cell divisions *in vitro* (**Figure 7C**).

Taken together, these results indicate that an individual stem cell’s response to Angpt1 treatment in culture depends upon its developmental history *in vivo*, with sensitivity to niche instruction being maximized in those cells that have divided 1-2 times previously. As has been suggested previously (Eldar & Elowitz 2010, Ridden et al. 2015), such an increase in stochasticity may provide a benefit to the stem cell population by increasing the range of division options available, thereby allowing it to ‘hedge its bets’ and remain primed to respond to different changes in circumstances (for example to replace tissue lost to injury) as required.

### Accumulation of HSCs with age requires individual cells to ‘remember’ their division history

To investigate our model further, we next sought to determine if it could be used to understand how stem cell numbers change with age. It has been widely observed that the frequency of immunophenotypic stem cells increases with age, yet this increase is associated with loss of regenerative potential and increase in linage bias (Pang et al. 2011, Sudo et al. 2000, Rossi et al. 2005, Morrison et al. 1996). Although thought to be a cell intrinsic phenomenon (Geiger et al. 2013), related to alterations in division patterns (Sudo et al. 2000), the precise underlying reason for these age-related changes are not yet well understood. To better understand this issue we used our model to predict how stem cell numbers would change, using the *in vivo* propensities for S-S, S-P and P-P divisions estimated from our PDC experiments (**Figure 7D**), and compared model predictions with experimental assessment of total LT-HSC numbers.

As in culture, we found that the estimated *in vivo* propensities for S-S, S-P and P-P divisions depended on the divisional history of the cell, with self-renewal potential declining with the number of previous divisions (**Figure 7D**). Similar to divisions *in vitro* the expansion factor Δ progressively declined with increasing number of prior divisions, indicating loss of self-renewal upon division, and cells that had divided two or three times previously were most stochastic in their division patterns (**Figures 7E-7F**). However, while culture was not able to support self-renewal for stem cells that had divided three or more times, we estimated that self-renewal ability is not lost until the fifth division *in vivo* (**Figure 7D**).

This observation has an important consequence for understanding proliferation with age: if each cell is able to divide five or more times then the stem cell pool as a whole will inevitably lose self-renewal potential, and will deplete beyond a critical age. To protect against this possibility requires aged stem cells (i.e those that have divided four times) to enter a dormant state, in which they do not proliferate under normal circumstances, and so are not removed from the system via further divisions. Our age-structured model of proliferation therefore takes into account the changing division propensities we observed, and assumes that stem cells enter a dormant state after four divisions. Full mathematical details of our proliferation model are given in the **Supplemental Material: Mathematical Modeling**.

To investigate this model we first monitored how total Evi1^+^ LT-HSC numbers changed with age in our mice. In accordance with previous reports we observed that LT-HSC numbers increased monotonically with age (**Figure 7G, left**, experimental data in black). Notably, we found that our model of stem cell proliferation – using the parameters independently estimated from PDC experiments – predicts this accumulation with excellent accuracy (**Figure 7G, left**), and estimates the HSC cell cycle time to be approximately 48 days (95% confidence interval [32, 67] days) in accordance with previous estimates (Cheshier et al. 1999). These results suggest that our model captures the essential dynamics of stem cell proliferation and aging.

Using our model we were also able to infer how the stem cell population structure changes with age. We found that successive waves of stem cells accumulate at distinct points of development (**Figure 7G, middle**): in young mice most stem cells have not divided, or have divided only once; through the middle of life the stem cell pool consists of a heterogeneous mix of cells that have divided between one and three times; while in old age the stem cell pool consists primarily of cells that have accumulated four divisions.

To validate our assumption of dormancy after four divisions we also compared how model predictions of stem cell numbers change if a fifth division is allowed. In this case, the stem cell pool is predicted to reach a maximum in mid life before declining into old age (**Figure 7G, right**, dotted lines). Thus, our model predicts that the accumulation of stem cell numbers with age is a consequence of history-dependent changes in the propensity of individual cells to self-renew, and entrance into a state of protective dormancy upon loss of self-renewal potential.

### The stem cell population reaches maximum potency in mid-life

Collectively, our data indicates that stem cell numbers increase with age, yet individual cells lose their regenerative potency with cell divisions. This suggests that a point of maximum *population* potency – in which stem cells are present in large numbers but have not yet lost self-renewal ability – should occur sometime in mid-life. To estimate this point we calculated how total renewing capacity, defined as the total number of stem cells multiplied by Δ, the expected number stem cells produced per stem cell division, changed with age under our model. We found that the point of maximum regenerative capacity occurred at approximately 135 days (19-20 weeks): at this point the expansion parameter Δ reached zero, indicating the onset of regenerative decline (**Figure 7H**), and stem cell dynamics became maximally stochastic (**Figure 7I**). From this point onward the regenerative capacity of the stem cell population decreases, divisions become more defined, and the observed increase in immunophenotypic stem cell numbers reflects the accumulation of a pool of aged stem cells with diminished self-renewal and proliferative capacity.

## DISCUSSION

There has been a long-running debate over stochastic versus instructive models of stem cell fate (Wagers et al. 2002, Suda et al. 1984, MacArthur et al. 2009, Ema et al. 2000, 2005, Ogawa 1999, Till et al. 1964). From the instructive perspective individual daughter cell identities are precisely determined by the niche, for example via cytokine instruction or juxtacrine signalling (Wu et al. 2007). In this view, while the output of cell divisions may vary, each division is precisely defined, albeit by mechanisms that may be hard to deconvolute. In contrast, in the stochastic perspective, daughter cells randomly adopt either a self-renewing or differentiated identity immediately after division – for example, due to fluctuations in the internal molecular state of the parent cell (Graf & Enver 2009). In this view, the niche does not directly instruct division dynamics *per se*; rather, it supports subsequent daughter cell development, for example to encourage maintenance of the self-renewing state or differentiation along a defined lineage. The crucial difference between these models is that in the stochastic view changes in cell identity arise from cell intrinsic stochastic processes that can be shaped by the niche (i.e. the niche is permissive but not directive), while in the instructive view such fluctuations may occur but are not functional, and changes in cell identity are determined by cellular responses to extrinsic signals that may be conditioned, but not driven, by intrinsic fluctuations.

There is evidence both for and against both hypotheses. In support of the instructive view, it has been shown that both self-renewal and lineage commitment of HSCs can be explicitly directed by extrinsic factors *in vitro*. Well known examples include the ability of stem cell factor (SCF) and Interleukin-3 (IL-3) to promote asymmetric HSC divisions (Takano et al. 2004), and macrophage colony stimulating factor (M-CSF) and granulocyte colony stimulating factor (G-CSF) to instruct fates of granulocyte-macrophage progenitors (Rieger et al. 2009). Similarly, the balance between symmetric and asymmetric HSC divisions may also be regulated *in vitro* by co-culture with different supportive cell populations (Wu et al. 2007). In support of the stochastic view it has been proposed that stochastic fluctuations in key lineage specifiers such as GATA1 and PU.1 – which regulate erythroid and myelomonocytic fate commitment respectively – prime individual stem cells for differentiation along defined lineages, with the niche serving to consolidate, rather than define, resulting cellular identities (Huang et al. 2007, Cross & Enver 1997, Akashi et al. 2003, Enver et al. 1998). The fact that mice lacking key lineage-specifying growth factors or HSCs lacking important growth factor receptors can still support healthy hematopoiesis (Lieschke et al. 1994, Stanley et al. 1994, Wu et al. 1995, Akashi et al. 1997, Lagasse & Weissman 1997) is indirectly supportive of this perspective. However, recent experiments using continuous long-term imaging of GATA1 and PU.1 expression in reporter cell lines has indicated that such fluctuations, while present, are not compatible with a model in which they initiate cell fate choice, but rather appear to consolidate prior fate commitment events (Hoppe et al. 2016). Collectively, these results suggest that the relationships between stochastic intracellular molecular states, niche instruction and stem cell fate commitment are complex and not completely described by current stochastic or instructive models.

Here, we propose a theoretical framework that reconciles these two perspectives. In our view, while the outcome of each division is inherently stochastic, depending on the random partitioning of intracellular resources, the outworking of this stochasticity is informed by instruction from the niche (here encoded by a simple threshold). Since it is the internal makeup of the cell relative to the expectation of the niche that determines the cell identity, this means that divisions may appear stochastic or instructive depending on the mismatch between niche expectation and internal state. So, for example, if the internal resource is particularly high in a cell (i.e. much higher than twice the niche threshold), then the stem cell identity is likely to be maintained through division independently of niche instruction. Similarly, if the internal resource is particularly low in a cell (i.e. only marginally higher than the niche threshold) then the stem cell identity is likely to be lost through division independently of niche instruction. Conversely, away from these two extremes (i.e. when the intracellular resource level is approximately twice the niche threshold) then the fate of the daughter cells will become sensitive to fluctuations around this threshold level, and will accordingly become sensitive to niche instruction. Thus, both inherent stochasticity and niche direction have important roles, with the balance between mechanisms depending on both intracellular makeup and extracellular context. Notably, because the internal state of the cell is determined by its resource reservoir, which dilutes continuously in the stem cell population as it proliferates, our framework views lineage commitment as a continuous process at the molecular level, that is discretized by the niche. This view is concordant with recent observations that individual HSCs gradually acquire lineage biases over time, rather than via a hierarchy of discrete transitions (Velten et al. 2017).

Our model is a simple conceptualization of an complex process, that is intended to facilitate exploration of the interplay between intrinsic and extrinsic regulatory processes, rather than a detailed molecular model. Nevertheless, it has a number of notable consequences. Firstly, within this framework, the propensity for an individual cell to self-renew depends upon its divisional history: cells that have never divided possess a large reservoir of internal resource and are therefore less likely to differentiate on division; while those that have divided numerous times previously have diminished intracellular resources and are more likely to lose stem cell status through division.

Secondly, assuming that stem cells do not divide synchronously, this history-dependence leads to the generation of an evolving age-structured population in which each stem cell in the pool exhibits a different propensity for self-renewal depending on its particular developmental history. In the **Supplemental Material: Mathematical Modeling** we discuss how stochasticity arising from variations in cell cycle times naturally generates a heterogeneous age-structured population. Importantly, this means that the potency of the population as a whole depends on the collective dynamics of an inherently heterogeneous mix of cells in which, due to their different divisional histories, no two stem cells are exactly alike.

Thirdly, the combination of loss of potency with division plus stochasticity in division dynamics means that when assessed in developmental time (i.e. mouse age) there is a transition from symmetric S-S divisions in young mice (in which HSCs have generally not yet divided many times and therefore retain an abundance of intracellular resources) to a mixture of S-S, S-P, and P-P divisions through mid life (as cells stochastically accumulate differing numbers of divisions and exhibit differing division propensities based on their individual division histories), and finally to predominantly P-P divisions in late life (when most HSCs have divided numerous times and therefore the resource has become critically depleted). At the population level this means that while division patterns are primarily deterministic in young and aged mice, they become highly stochastic through the middle of life, and more sensitive to the resource threshold imposed by the niche.

Fourthly, our model predicts that in order to maintain an active stem cell pool with age, individual stem cells must be able to ‘count’ their divisions and enter a dormant state after their fourth division. This prediction is notable because it is in accordance with recent work, based upon the dynamics of label retaining HSCs with age, which also predicted that HSCs enter a dormant state after four divisions (Bernitz et al. 2016). We currently know very little about how such memory processes work. However, they may be inherent to regulating cellular dynamics with age, and therefore be important in understanding age-related regenerative decline. We anticipate that further investigation of cellular memories – for example using label retention dynamics or genetic labelling strategies in combination with single cell molecular profiling – will be fruitful avenues for future work.

Although the model that we propose describes the data well, a number of unresolved questions remain. Most notably, while our model is agnostic regarding the exact nature of the intracellular resource(s), their precise nature is clearly of importance. Candidate factors should be highly expressed in juvenile HSCs, yet lose expression with age, and be expressed at low levels in aged/dormant HSCs. There are a number of possibilities. Dormant stem cells are known to express low levels of cell cycle regulators such as Cdk6 (Cabezas-Wallscheid et al. 2017, Laurenti et al. 2008) and Myc (Laurenti et al. 2008), and depend on glycolysis for energy production in order to maintain low levels of reactive oxygen species levels (Ito & Suda 2014). Conversely, key DNA- and RNA-binding proteins such as Sox17 and Lin28 are known to be expressed in juvenile, but not adult HSCs (He et al. 2011, Kim et al. 2007, Copley et al. 2013, Yuan et al. 2012). Interestingly, forced re-expression of these factors in adult stem and progenitor cells is able to restore an immature phenotype, suggesting that changes in cellular identities following the loss of key intracellular resources can be reversed (He et al. 2011, Yuan et al. 2012). It is likely that our putative resource consists of a cocktail of important intracellular determinants, and it will be interesting to see if different factors have different dilution dynamics through divisions, and how combinatorial changes in expression patterns that arise due to differential dilution dynamics affect cellular responses to the niche.

Although our model assumes a single *in vivo* niche threshold for convenience (and to account for stem cell regulation under homeostatic conditions) it is likely that niche instruction can change, and the balance between instructive and stochastic mechanisms may itself be regulated and/or evolve with age. In this case, the balance between the inherent stochasticity of intracellular molecular dynamics and extracellular regulation of cell identities by the niche may transiently alter to expand stem cell numbers if required, for example under conditions of stress. An interesting possibility is that cells that have previously ‘differentiated’ under weakly instructive niche conditions may retain the capacity to spontaneously revert back to a stem cell status if placed in a strongly instructive niche or in conditions that enhance the production of intracelluar resources. It has been argued that such ‘dynamic heterogeneity’ is a robust strategy for maintaining homeostatic stem cell activity (Greulich & Simons 2016); if so, this could be an important feature of stem cell dynamics that is, as yet, poorly understood and worth further investigation. Similarly, it may also be possible that dormant stem cells can be reactivated, and re-enter the regenerative pool; indeed such transiting of HSCs in and out of dormant and activated states in response to stress has been observed (Wilson et al. 2008). The fact that HSCs enter a dormant state after four divisions may therefore be a protective mechanism that ensures that individual cells do not lose the latent capacity to be reactivated, and therefore preserves a pool of stem cells that can respond in times of stress. We anticipate that investigation of the extent to which such phenotypic switching occurs at the single cell level, for example using high-resolution single cell tracking in combination with high-throughput single cell sequencing, will be a fruitful direction for future work.

In summary, our work indicates that stochastic and instructive regulatory mechanisms work in harmony to regulate the activity of individual stem cells and produce a heterogeneous, age-structured stem cell population that is able to maintain healthy life-long hematopoiesis without proliferative exhaustion.

## ACKNOWLEDGEMENTS

We would like to thank Dr. Mineo Kurokawa (The University of Tokyo, Japan) for providing Evi1-GFP knockin mice. We also thank Dr. Gou Young Koh (KAIST, Korea) for providing recombinant Angpt1 protein. This research was funded by the funding program for Next Generation World-Leading Researchers (NEXT Program, grant number LS108), Scientific Research (B) (General), grant number 17H04208, from the Ministry of Education, Culture, Sports, Science and Technology (MEXT) of Japan, Takeda Science Foundation, the Medical Research Council, United Kingdom, grant number MC PC 15078, and a donation from Fujino Brain Research, Ltd.

## AUTHOR CONTRIBUTIONS

Conceptualization BDM, FA, TS; Methodology BDM, FA; Software BDM, PSS; Formal Analysis BDM, FA, PSS; Investigation, AR, BDM, FA, KH, YMI; Resources MPL; Data Curation BDM, PSS; Writing - Origninal Draft BDM, FA; Writing - Reviewing & Editing BDM, FA, MPL, PSS, TS; Visualization, BDM, PSS; Supervision BDM, FA, MPL, TS; Project Administration, BDM, FA; Funding Acquisition BDM, FA.

## DECLARATION OF INTERESTS

The authors declare no competing interests.

## STAR METHODS

### CONTACT FOR REAGENT RESOURCE SHARING

Further information an requests for resources should be directed and will be fulfilled by the Lead Contact, Ben MacArthur.

### EXPERIMENTAL MODEL AND SUBJECT DETAILS

#### Mice and cells

C57BL/6 (B6-Ly5.2), C57BL/6 mice congenic for the Ly5 locus (B6-Ly5.1) were purchased from Sankyo-Lab Service (Tsukuba, Japan). Evi1-GFP knock-in mice were provided by Dr. Kurokawa (The University of Tokyo, Japan). Mice expressing Cre recombinase under the control of the Type I collagen (Col1a1) promoter (Col1a1-Cre) and inducible Angpt1 transgenic (CAGp-IND-COMP-Angpt1 Tg) mice (Hato et al. 2009) were used to obtain constant and specific expression of Angpt1 in osteoblasts (Col1a1-Cre(+): Angpt1 Tg). The gene recombination experiment safety and animal experiment committees at Keio and Kyushu Universities approved this study, and all experiments were carried out in accordance with the guidelines for animal and recombinant DNA experiments at both Universities.

### METHOD DETAILS

#### Flow cytometry and antibodies

The following monoclonal antibodies (Abs) were used for flow cytometry and cell sorting: anti-c-Kit (2B8, BD Biosciences, 1:100), -Sca-1 (E13-161.7, BD Biosciences, 1:100), -CD4 (RM4-5, BD Biosciences, 1:100), -CD8 (53-6.7, BD Biosciences, 1:100), -B220 (RA3-6B2, BD Biosciences, 1:100), -TER-119 (BD Biosciences, 1:100), -Gr-1 (RB6-8C5, BD Biosciences, 1:100), -Mac-1 (M1/70, BD Biosciences, 1:100), -CD41 (MWReg30, eBioscience, 1:100), -CD48 (HM48-1, Biolegend, 1:100), -CD150 (TC15-12F12.2, Biolegend, 1:100), -CD34 (RAM34, BD Biosciences, 1:20), -CD45.1 (A20, BD Biosciences, 1:100), -CD45.2 (104, BD Biosciences, 1:100), and -Tie2 (TEK4, eBioscience, 1:100). A mixture of CD4, CD8, B220, TER-119, Mac-1, and Gr-1 was used as the lineage mix. To isolate Evi1^+^ LT-HSCs, Evi1-GFP mouse bone marrow (BM) cells were purified using a two-step protocol. First, LT-HSCs were enriched by positive selection for c-Kit expression using anti-CD117 immunomagnetic beads (130-091-224, Miltenyi Biotec Inc. 1:5 dilution). Second, c-Kit^+^ enriched cells were stained with a set of fluorophore-conjugated antibodies (lineage mix, anti-Sca-1, anti-c-Kit, anti-CD48, anti-CD150, and anti-CD34). For the analysis of Tie2 expression, BM cells from Evi1-GFP mice were stained with the lineage mix, anti-Sca-1, anti-c-Kit, anti-CD48, anti-CD150, anti-CD34, and anti-Tie2 antibodies. Stained cells were analyzed and sorted by flow-cytometry using a FACS Aria III (BD Biosciences).

#### Microarrays

LSKCD41^−^CD48^−^CD150^+^CD34^+^ cells, LSKCD41^−^CD48^−^CD150^+^CD34^−^ Evi1-GFP^−^ cells, and LSKCD41^−^CD48^−^CD150^+^CD34^−^Evi1^+^ cells were selected sorted from 8-week old mice, and RNA was isolated from each population using an RNeasy Plus Micro Kit (Qiagen) according to the manufacturer’s instructions. RNA quality was confirmed with a 2100 Bioanalyzer (Agilent Technologies, Santa Clara, CA). Total RNA (1.5ng) was reverse transcribed and amplified using the Ovation Pico WTA System V2 (NuGEN) with Ribo-SPIA technology. Amplified cDNA (5*µ*g) was fragmented and labeled with biotin using Encore Biotin Module (NuGEN), and hybridized to a GeneChip Mouse Genome 430 2.0 Array (Affymetrix) according to the manufacturer’s instructions. All hybridized microarrays were scanned using a GeneChip Scanner (Affymetrix). Relative hybridization intensities and background hybridization values were calculated using the Affymetrix Expression Console software (Affymetrix). Raw signal intensities for each probe were calculated from hybridization intensities. Raw signal intensities were normalized using the MAS5 method.

#### Paired daughter cell assay

Individual Evi1^+^ LT-HSCs were selected from 4-week old, 8-week old, and 18-month old mice and directly sorted (one-per-well) into fibronectin-coated 96 well U-bottom plates. Cells were cultured in SF-O3 medium in the presence of 0.1% BSA, 100 ng/ml SCF, and 100 ng/ml TPO with or without Angpt1 (1 *µ*g/ml, a gift from Dr. Koh, Korea Advanced Institute of Science and Technology). After 2 days of culture, daughter cells derived from single Evi1^+^ LT-HSCs were separated using a micro-manipulator (NARISHIGE, Japan) and transferred one-per-well to PCR plates containing a mixture of CellsDirect 2× Reaction Mix (Invitrogen), pooled 0.2× TaqMan Gene Expression Assays, and SuperScript III RT/Platinum Taq Mix (Invitrogen). Reverse transcription (RT) (at 50°C for 15 min), an initial denature step (95°C for 2min), and specific target amplification (STA) (22 cycles of 15s at 95°C alternating with 1 min at 60°C) were performed sequentially. Pre-amplified cDNA was diluted with TE buffer (1:5). Gene expression read-out was performed using BioMark 96*·*96 Dynamic Array and BioMark system (Fluidigm) according to manufacturer’s instructions. TaqMan Gene Expression Assays used in this study are listed in **Supplemental Table S1**.

#### Training data for classifier

To prepare the training data needed for our artificial neural network classifier, we profiled the expression of the same set of 96 HSC-related genes that we used in the PDC assay (see **Supplemental Table S1**) in 96 individual cells isolated without culture from the following nine populations:

1. LSKCD41^+^CD48^+^CD150^+^ progenitor cells taken from 4-week old mice (designated week 4 CD150^+^)
2. LSKCD41^+^CD48^+^CD150^−^ progenitor cells taken from 4-week old mice (designated week 4 CD150^−^)
3. LSKCD41^−^CD48^−^CD150^+^ Evi1^+^ stem cells taken from 4-week old mice (designated week 4 Evi1^+^)
4. LSKCD41^−^CD48^−^CD150^−^CD34^+^ progenitor cells taken from 8-week old mice (designated week 8 CD150^−^)
5. LSK progenitor cells taken from 8-week old mice (designated week 8 LSK)
6. LSKCD41^−^CD48^−^CD150^+^CD34^−^ Evi1^+^ taken from 8-week old mice (designated week 8 Evi1^+^)
7. LSKCD41^−^CD48^−^CD150^−^CD34^+^ progenitor cells taken from 18-month old mice (designated 18 month CD150^−^)
8. LSKCD41^+^CD48^+^CD150^+^CD34^+^ progenitor cells taken from 18-month old mice (designated 18 month CD150^+^)
9. LSKCD41^−^CD48^−^CD150^+^CD34^−^ Evi1^+^ stem cells taken from 18-month old mice (designated 18 month Evi1^+^)

Individual cells from each population were sorted directly into a mixture of CellsDirect 2× Reaction Mix, pooled 0.2× TaqMan Gene Expression Assays, and SuperScript III RT/Platinum Taq Mix. Following RT-STA (described above), gene expression was then analyzed using BioMark 96*·*96 Dynamic Arrays (Fluidigm).

#### Bone marrow transplantation

To evaluate the number of LT-HSCs, Ly5.1^+^ LSKCD41^−^CD48^−^CD150^+^CD34^−^Evi1-GFP^−^ and -Evi1^+^ cells were transplanted into lethally irradiated Ly5.2^+^ mice using 2 × 10^5^ competitor cells. The percentage of donor-derived cells in PB was analyzed monthly by flow cytometry. To evaluate the effect of the Angpt1 on the regulation of LT-HSC number in vivo, LSKCD48^−^ cells were isolated from Col1a1-Cre(+);Angpt1(+) Tg mice or Col1a1-Cre(–);Angpt1(+) control mice and the limiting number of cells (20, 30, 40, and 50 cells for Angpt1 Tg mice, 50, 75, and 100 cells for control mice) were transplanted into lethally irradiated recipient mice with 2 × 10^5^ competitor cells. The percentages of donor-derived cells were analyzed 4 months after BMT. PB chimerism of > 1.0% at 16 weeks after BMT was taken to verify long-term engraftment. To investigate the effect of Angpt1 in the maintenance of LT-HSC number *in vitro*, PDCs derived from the control culture or Angpt1 supplemented culture were transplanted into lethally irradiated recipient mice with 2 × 10^5^ competitor cells (1, 2, 3, 4, and 6 pairs of PDCs from each culture condition). The percentages of donor-derived cells were analyzed 4 months after BMT. PB chimerism of > 0.3% at 16 weeks after BMT was taken to verify long-term engraftment. The frequency of LT-HSCs and statistical significance were determined using the ELDA software (Hu & Smyth 2009).

### Manufacture of poly(ethylene glycol) (PEG) microwells

PEG-based hydrogel microwells were manufactured as previosuly described (Roch et al. 2017). Briefly, arrays of hydrogel microwells were fabricated using standard photolithography to etch the microwell pattern into a silicon substrate onto which liquid thermocurable poly(dimethylsiloxane) (PDMS) was polymerized. This template was then used to cast hydrophilic polymer precursors of PEG that were covalently crosslinked to form inert hydrogels. PEG hydrogel microwell arrays were then removed from the PDMS, equilibrated in PBS, and attached to the bottom of tissue culture wells.

## QUANTIFICATION AND STATISTICAL ANALYSIS

### Microarray data analysis

Differentially expressed genes from microarray studies were determined using the R package limma (Ritchie et al. 2015). Genes withe FDR corrected p-values < 0.05 (see **Supplemental Table S2**) were used for gene set enrichment analysis using the biological processes Gene Ontology (GO) in DAVID (Huang et al. 2008).

### Normalization and analysis of single-cell qRT-PCR data

Cycling threshold (CT) values ≥ 28 were considered absent. Raw CT values were normalized using the median CT values of loading controls (Gapdh) for each array. Normalised CT values were then transformed linearly to expression threshold (ET) values ranging from 0 (absent) to 28 (maximum expression). Gene expression asymmetry for each given gene was assessed by comparing ET levels between daughter cells. Specifically, if *g*_*i j*_*i* denotes the ET level of gene *g*_*i*_ in daughter cell *j* = 1, 2 then expression asymmetry is given by *a*_*i*_ = |*g*_*i*1_ *-g*_*i*2_|. Total asymmetry for the division was then defined as Σ_*i*_ *a*_*i*_.

### Clustering and dimensionality reduction

All clustering and dimensionality reduction was performed in R (version 3.1.2 or later) and MATLAB^®^ (The MathWorks Inc., Natick, MA, 2016, version 8.5 or later) using standard routines. Hierarchical clustering was performed in R using Euclidean distance and average linkage, t-distributed stochastic neighbour embedding (t-SNE) was performed in MATLAB^®^ using the Toolbox for Dimensionality Reduction (Van Der Maaten et al. 2009) with default settings.

### Estimation of dispersion

To the *i*th cell in the population we associate a gene expression vector *G*_*i*_ = (*g*_1_, *g*_2_, *…, g*_96_) ∈ℝ^96^, which records its expression status with respect to the 96 genes we measured. Assuming that there are *n* cells in the population, the mediancentre is that point *M* = (*m*_1_, *m*_2_, *…, m*_96_) ∈ℝ^96^ such that 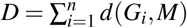 is minimum, where 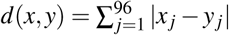 is the *L*_1_-distance. The mediancentre is a multivariate generalization of the univariate median (Gower 1974). The dispersion of each cell is its distance to mediancentre *d*(*G*_*i*_, *M*), and the dispersion of the population is the minimized value of *D*. The dispersion is a simple statistic that can be used in hypothesis testing to compare the multivariate variability in different populations (Van Valen 2005).

### Artificial neural network

We used a deep feedforward artificial neural network (ANN) to distinguish between the nine training samples, and thereby identify the regenerative status of the cell (stem or progenitor cell) and its age (4-week, 8-week or 18-month old) directly from its transcriptional profile. Network parameters were optimized using grid search over a range of network architectures, including number of hidden layers and nodes per layer, activation functions and regularization regimes. We found that the ANN constructed with two hidden layers of 150 rectifying linear units (ReLUs) each, trained with cross-entropy loss function using dropout (20% dropout for input and 50% and 75% for the two hidden layers respectively) and additional *L*_1_ and *L*_2_ regularization (both with 0.002 penalty), with soft-max output performed best. For each set of network parameters optimization was performed with 10-fold cross-validation using stochastic gradient descent and back-propagation over 1,000,000 epochs. All reported metrics refer to cross-validated results. All training and optimization was performed in R using the H2O deep learning library (Cook 2016).

### Model fitting

Model parameters were estimated by minimizing the residual sum of squares between the experimental data and the model. Optimization was performed using the Levenberg-Marquardt algorithm, implemented in MATLAB^®^ (The MathWorks Inc., Natick, MA, 2016, version 8.5 or later) as part of the Optimization Toolbox. The full model has the following parameters:

*n* = Resource copy number in founder cell *in vivo*,

*q* = Resource replenishment rate *in vivo*,

*T*_vivo_ = Niche threshold *in vivo*

*k* = Shape parameter for Gamma distribution describing cell cycle time distribution *in vivo*,

*θ* = Scale parameter for Gamma distribution describing cell cycle time distribution *in vivo*,

*α* = Resource asymmetry *in vitro*,

*T* = Niche threshold *in vitro*.

All *in vivo* parameters refer to the developmental history of stem cell population before the experiment started and are therefore common to all PDC experiments. When fitting, we restricted parameter bounds to ensure that experiments conducted in different culture conditions gave estimates of these parameters that were statistically equivalent (i.e. within 95% confidence intervals, see (Bates & Watts 1988)). To reduce the number of free parameters further we made the following additional assumptions: (1) since it is the threshold *T*_vivo_ relative to the resource copy number in the founder cells *n* that defines the dilution dynamics, we set *n* = 1, 000 and fitted for *T*_vivo_ as a free parameter. Similar results may be obtained by varying *n* and scaling *T*_vivo_ accordingly. (2) To allow asymmetric divisions *in vitro* that mimic dilution dynamics *in vivo* we set 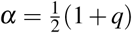. To describe how LT-HSC numbers changed with age we imported estimates of *in vivo* parameters from PDC experiments into our model of age-structured proliferation (see **Supplemental Material: Mathematical Modelling**). This model has one additional parameter *M*(0), the founder stem cell number, which was then inferred by fitting to the experimentally observed data as above.

## DATA AND SOFTWARE AVAILABILITY

### Fluidigm Dynamic Arrays

Single-cell gene expression data reported in this paper is available from ArrayExpress under accession number E-MTAB-7504.

### Microarrays

Ensemble-cell microarray data reported in this paper is available from ArrayExpress under accession number E-MTAB-7321.

## Supplemental Material

### Mathematical Modeling

#### Stochastic model of proliferation and the expansion factor Δ

The simplest stochastic model of stem cell proliferation assumes the following set of events:

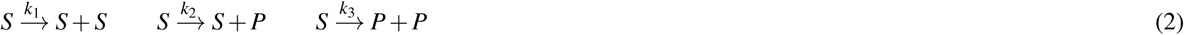

where *S* denotes a stem cell and *P* denotes a progenitor. The first event is a symmetric S-S division, in which 2 stem cells are produced via division, resulting in a gain of one stem cell to the pool; the second is an asymmetric S-P division, in which one stem cell is produced via division, resulting in no change to the stem cell pool; and the third is a symmetric P-P division, resulting in a loss of one stem cell from the pool. Let *p*_*n*_ be the probability that there are *n* stem cells in the pool at time *t*. The master equation describing these dynamics is:

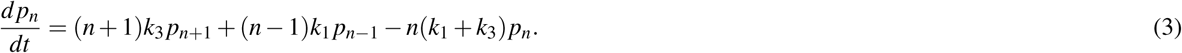

Let *M*(*t*) = Σ_*n*_ *np*_*n*_ denote the expected number of stem cells in the pool at time *t*. The equation for *M*(*t*) may easily be found to be (Van Kampen 1992):

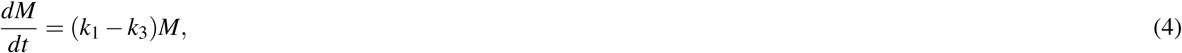

from which we obtain, *M* = *M*(0)exp([*k*_1_ - *k*_3_]*t*). Assuming that the cell cycle time is constant, we may write *k*_1_ = *pk* and *k*_3_ = *qk*, where log(2)*/k* is the cell cycle time, *p* is the probability that an S-S division occurs and *q* is the probability that a P-P division occurs. Thus, *M* = *M*(0)exp(*k*Δ*t*), where Δ = *p-q*. If Δ > 0 then S-S divisions are more likely than P-P divisions and the stem cell pool will expand, while if Δ < 0 then P-P divisions are more likely than S-S divisions and the stem cell pool will deplete. If Δ = 0 then S-S and P-P divisions are equiprobable, and homeostasis is maintained (i.e. *M* = *M*(0) for all *t*).

#### Resource dynamics: general mathematical framework

We assume that each long-term repopulating hematopoietic stem cell (HSC) contains a supply of a molecular resource *X*. Let *x*_0_ denote the copy number of *X* in a stem cell and assume that immediately prior to cell division *x*_0_ = *n*. Upon division this resource is partitioned between the two daughter cells. Let *x*_*i*_ denote the copy numbers of *X* in daughter *i* = 1, 2, and let *f*_*i*_(*k*) be the corresponding probability that daughter *i* = 1, 2 inherits *k* copies of *X*.

Now, assuming that the total amount of *X* does not change through the division, if daughter 1 inherits *k* copies of *X* then daughter 2 necessarily inherits *n-k* copies. Thus,

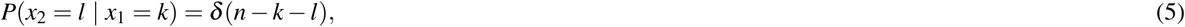

where *δ* is the Kronecker-delta function. The joint distribution *f* (*k, l*) = *P*(*x*_1_ = *k, x*_2_ = *l*) for expression of *X* in the pair of daughter cells may then be found via Bayes’ theorem,

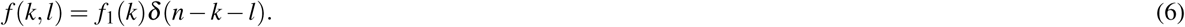

To evaluate this expression a little care must be taken because, although it is mathematically convenient to introduce an ordering for the two daughter cells, this order is not observed experimentally, and it is the *observed* statistics that we are concerned with. Experimentally, cells are observed in pairs and assigned labels (daughter 1 or 2) randomly and uniformly. To take account of this we introduce the ‘observed’ marginal probability, 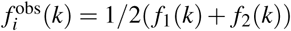 for *i* = 1, 2. Thus, the statistics of cells designated ‘daughter 1’ are indistinguishable from those designated ‘daughter 2’, i.e. the order in which the daughter cells are observed does not affect their identity.

A simple model of cell division is to assume that each copy of *X* is retained independently in daughter 1 with constant probability *α*. In this case *x*_1_ is binomially distributed, and

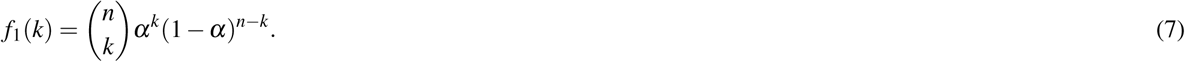

Thus, if *α* = 1*/*2 then the division is perfectly symmetric at the molecular level (on average both daughters inherit the same amount of the resource *X)* while if *α* = 0, 1 then the division is perfectly asymmetric at the molecular level (one daughter inherits all the resource, and the other none of it).

In practice it is not likely that the identity of an individual cell is determined by the copy number of a single molecular resource (here *X)*. Rather, cellular identities are determined by complex co-expression patterns of a variety of different factors, which may be partitioned in regulated way. However, the argument above still applies to each regulatory factor when considered separately.

The above argument concerns expression of the resource *X* in daughter cells. However, the copy number of these resources are not directly observed in experiment. Rather we assign an identity to each daughter based upon its gene expression pattern using an artificial neural network. This method establishes a map from gene expression status of the cell to its cellular identity. In the case that cells are classified as either stem or progenitor cells, and assuming that the expression of *m* genes are assessed in each cell, this results in a map:

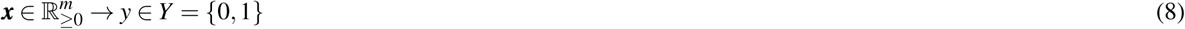

where *y* = 0 denotes a progenitor cell identity and *y* = 1 denotes a stem cell identity. To understand the experimentally observed paired daughter cell patterns it is therefore necessary to have a model of how cellular identities arise from molecular states.

Let *ρ*_*k*_ be the probability that a daughter cell with *k* copies of *X* after cell division adopts the stem cell identity. From Bayes theorem we may obtain the probability *p*_*i*_ that daughter *i* adopts a stem cell identity

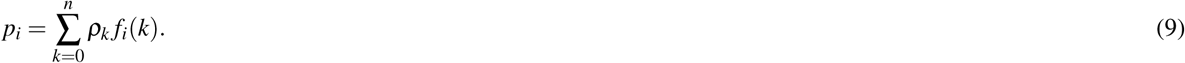

Since we do not observe *f*_*i*_(*k*) some care should again be taken: daughter cells are observed in pairs and assigned identities (daughter 1 or 2) randomly and uniformly, so the probability of observing a stem cell is independent of daughter index, and is given by

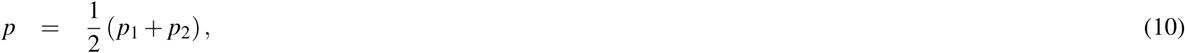

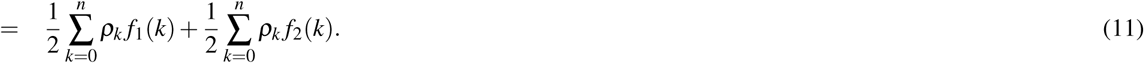

Furthermore, since *P*(*x*_2_ = *k* | *x*_1_ = *k*′) = *δ* (*n-k-k*′) it is immediate that

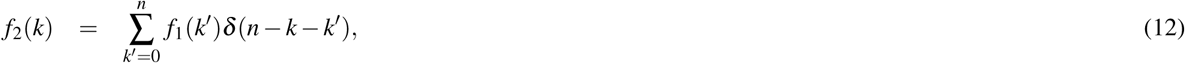

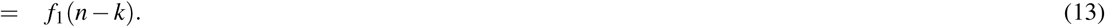

Thus, without loss of generality we can represent the dynamics in terms of *f*_1_(*k*). For simplicity of notation we will denote *f*_1_(*k*) = *f* (*k*). The probability *p* of observing a stem cell may then be written

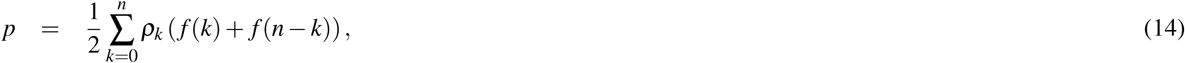

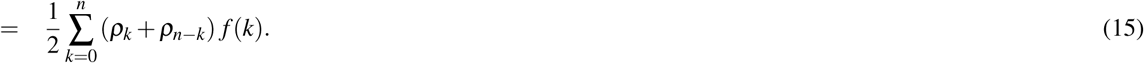

Denoting the identity of daughter *i* = 1, 2 by *y*_*i*_ ∈ *Y*, the joint probability *p*_*ab*_ = *p*(*y*_1_ = *a, y*_2_ = *b*) is a bivariate Bernoulli distribution, which may be written in general form as

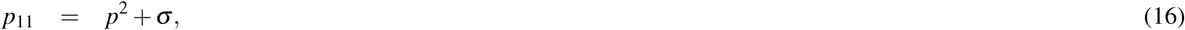

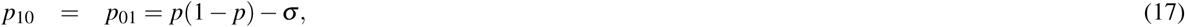

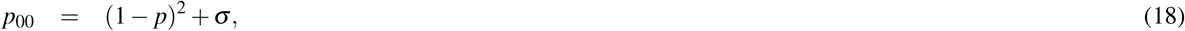

where *σ* is the observed covariance between the daughter identities, and *p*_11_ = 𝔼 (*y*_1_*y*_2_) is given by:

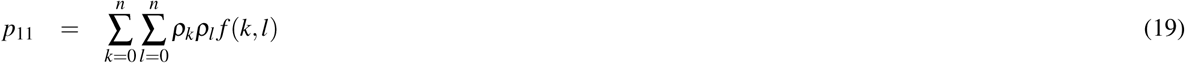

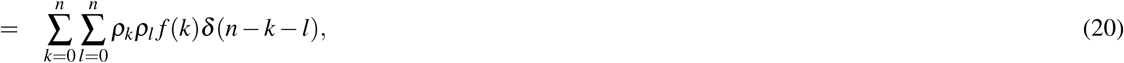

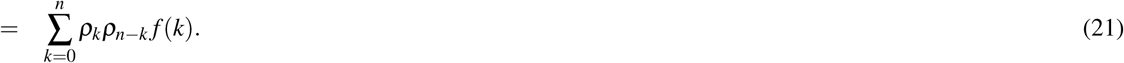

The observed covariance between classified daughters may then be obtained in terms of *ρ*_*k*_ and *f* (*k*), which quantify how the resource *X* is partitioned through cell division, and the propensity of the daughter cells to differentiate given their expression of *X* respectively. In particular,

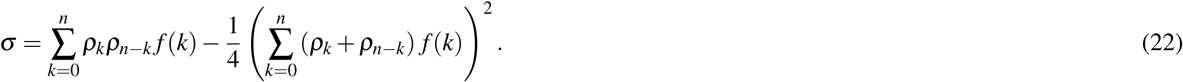

The probability *p* that a randomly chosen daughter cell is a stem cell and the covariance between daughter cells *σ*, given by Eqs.(15) and (22) respectively, fully determine the general form of the model.

#### Stem cell aging: a resource partitioning model

To put this general model in context it is helpful to consider a specific instance. As a simple model of differentiation we assume that each daughter cell adopts a stem cell identity if it inherits enough resource, and differentiates if it inherits too little resource. Thus, *ρ*_*k*_ = 1 if *k* > *T* and *ρ*_*k*_ = 0 if *k* ≤ *T* for some threshold *T*. In this case *ρ*_*k*_*ρ*_*n-k*_ = 1 when *k* > *T* and *k* < *n-T* and is zero otherwise. These two inequalities can only both be satisfied if *T* < *n-T*, or alternatively if *T* < *n/*2. Thus *ρ*_*k*_*ρ*_*n-k*_ = 0 for all *k* if *T* ≥ *n/*2. In this case, *p*_11_ = 0, and *σ* = *-p*^2^ ≤ 0. Alternatively, if *T* < *n/*2, then

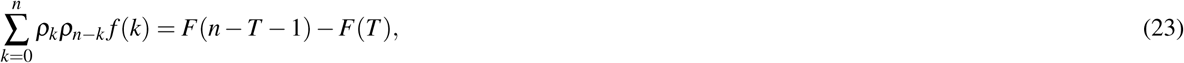

where *F*(*k*) is the cumulative distribution function for *f* (*k*). Similarly,

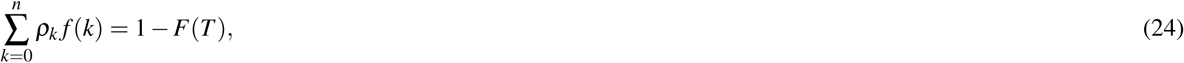

and

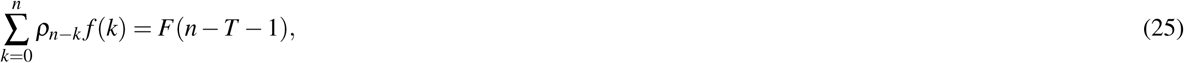

and therefore,

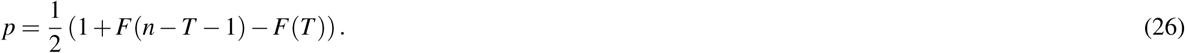

Thus, for *T* < *n/*2

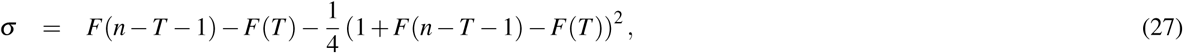

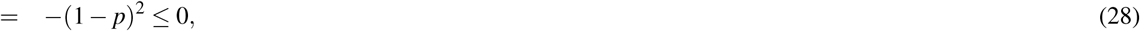

In this case,

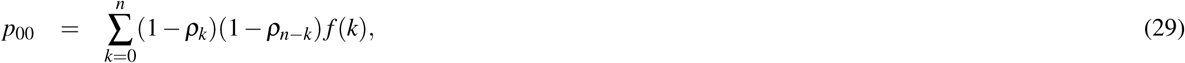

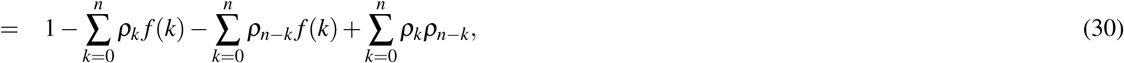

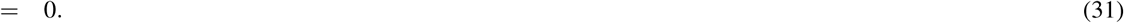

In summary, if *T* ≥ *n/*2 then

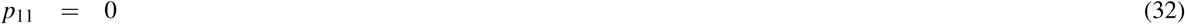

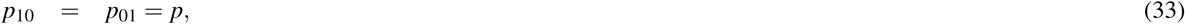

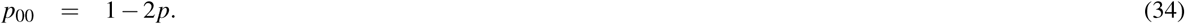

Similarly, if *T* < *n/*2 then

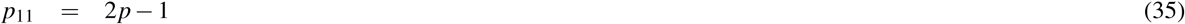

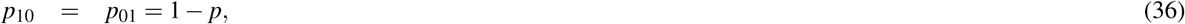

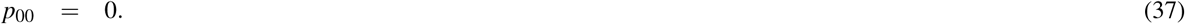

From Eq.(26) note that if *T* < *n/*2 then *F*(*n-T)* > *F*(*T)* and therefore *p* > 1*/*2, while if *T* ≥ *n/*2 then *F*(*n-T)* ≤ *F*(*T)* and *p* ≤ 1*/*2, so these probabilities are well defined. In the special case that *T* = *n/*2, *p* = 1*/*2 and therefore *p*_00_ = *p*_11_ = 0, and *p*_10_ = *p*_01_ = 1*/*2. In this case, only asymmetric cell divisions occur and this simple model gives a mechanistic basis for the standard model of cellular differentiation.

#### Evolution of stem cell dynamics with age

So far we have assumed that the resource *X* is present in the parent cell at a fixed level. To understand how stem cell division dynamics evolve with age we need to have a model of how expression of *X* varies in the stem cell pool with age.

Consider a single founder cell in which *x*_0_ = *n*. Let this cell divide successively to produce daughter cells, then granddaughter cells etc. We wish to know the probability distribution of the expression of *X* in the descendant population at the *g*-th generation. Let *α*_*i*_ be the probability that daughter cell 1 inherits each copy of *X* at generation *i*. In general these division probabilities may differ from each other, allowing for partitioning dynamics to change with cell division history. However, because cell divisions *in vivo* have been observed to be primarily symmetric (Bernitz et al. 2016), we assume that *α*_*i*_ = 1*/*2 for all generations (i.e. each division id symmetric and identical to the last at the *molecular* level). This assumption reduces model parameters, and so helps avoid over fitting, and provides a model that is analytically tractable. Under these dynamics the expected copy number of the resource in a cell that has divided *g* times is *n/*2^*g*^. Thus, the resource rapidly dilutes, unless there is some mechanism to replenish it between divisions. To account for resource replenishment, we assume that each copy of *X* that is ‘lost’ to the sister cell is recovered with probability *q* before the next cell division. Thus, the dilution dynamics are governed by *q*. In particular, the expected copy number of *X* in a cell that has divided *g* times previously is then 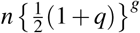.

The dynamics of resource dilution *in vivo* are then governed by a Galton-Watson branching process (Van Kampen 1992) and the probability *p*_*k*_(*g*) that a randomly selected stem cell from the pool contains *k* copies of *X* after the *g*-th division is

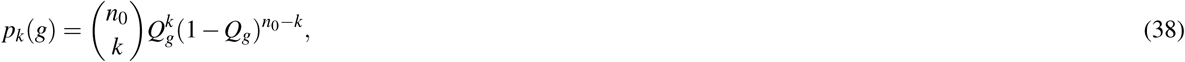

where 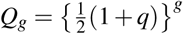.

#### Division dynamics in vitro

The model above accounts for resource dilution dynamics *in vivo*. To investigate stem cell divisions *in vitro*, we assume that differentiation occurs if the copy number of *X* falls below a critical threshold *T* > *T*_vivo_ that is determined by the culture conditions. To allow for stress-induced changes to divisions patterns we also assume that daughter cell 1 inherits each copy of *X* with probability *α* (which may be different from 1*/*2, thereby allowing asymmetric divisions at the molecular level) and assume that the daughter cells are harvested before the resource has time to replenish.

Under these assumptions we need to consider three possibilities based upon the copy number *k* of *X* in the parent cell: (1) *k* ≤ *T*_vivo_, in which case the parent cell has differentiated; (2) *T* < *k* ≤ 2*T*, in which case P-P and S-P divisions may occur, but there is insufficient resource in the parent cell to allow both daughter cells to inherit enough resource to allow an S-S division; (3) *k* > 2*T*, in which case S-S and S-P divisions may occur, but there is too much resource to allow a P-P division. Thus, the probabilities *p*_11_(*g*), and *p*_00_(*g*) that a stem cell that has divided *g* times *in vivo* will perform an S-S or P-P division *in vitro* respectively are:

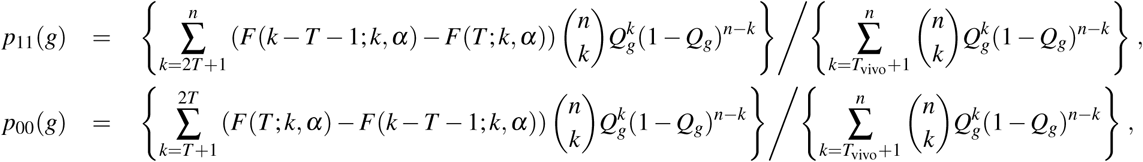

#### Conversion from cell division generation to developmental time

The above results quantify the self-renewal potential of a stem cell in terms of the number of times it has divided previously. By doing so we are measuring the ‘age’ of the cell in terms of the number of times it has divided. However, this developmental age is not amenable to experimental assessment. Rather, experiments are conducted in which stem cells are selected from mice of different natural ages. We therefore need a model of how cell intrinsic time (i.e. the number of times a given cell has divided) relates to developmental time. We must thus answer the question: if a cell is selected randomly from the HSC pool in a mouse of age *t*, what is the probability that the selected cell has divided exactly *r* times previously?

To address this question requires a statistical model of cell proliferation. It has previously been observed that the cell cycle may be modeled as a sequence of independent Poisson processes, and the cell cycle time is consequently well described by a Gamma distribution (Yates et al. 2017). Let *s*_*r*_ denote the length of time taken for a randomly selected stem cell to complete its *g*-th division (i.e. the time between completion of the (*g-* 1)th division and the beginning of the *g*-th division). Assuming that each cell cycle is independent of the last, and is modelled by a sequence of Poisson processes, we obtain *s*_*g*_ ∼ Gamma(*k*_*g*_, *θ)*, where *k*_*g*_ and *θ* determine the shape and scale of the Gamma distribution respectively. Since the sum of independent Gamma distributed random variables of equal scale is also Gamma distributed, the random variable *τ*_*g*_ describing the total time taken from the beginning of the first cell division the end of the *g*-th division is then distributed according to *τ*_*r*_ ∼ Gamma(*K*_*g*_, *θ)*, where 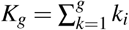.

Let *p*(0, *t*) be the probability that a randomly selected cell has not divided at time *t*. In this case,

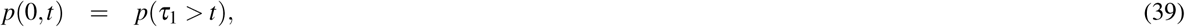

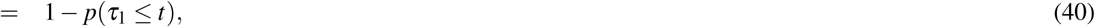

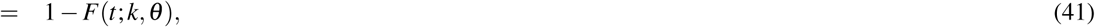

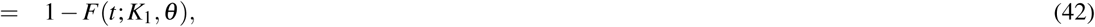

where *F*(*t*; *k, θ)* = *F*(*t*; *K*_1_, *θ)* is the cumulative distribution function for Gamma(*k, θ)*. Similarly, if *p*(1, *t*) is the probability that a randomly selected cell from the HSC pool in a mouse of age *t* has divided exactly once previously, then:

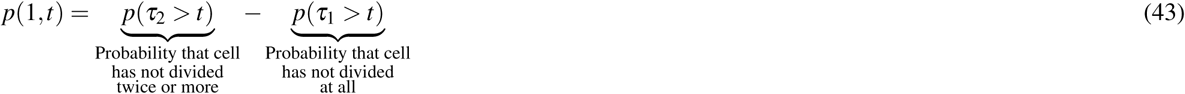

from which we obtain

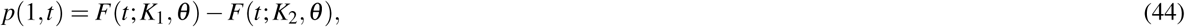

where *F*(*t*; *K*_*g*_, *θ)* is the cumulative distribution function for Gamma(*K*_*g*_, *θ)*. A similar argument extends to higher probabilities and in general,

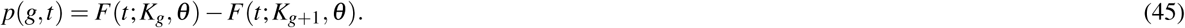

These probabilities allow us to translate from cell intrinsic time (i.e. number of times a cell has divided) which is not observable, to developmental time (i.e. mouse age) which is. Thus, the probabilities *p*_11_(*t*) and *p*_00_(*t*) of observing an S-S or P-P division respectively from a stem cell selected from a mouse of age *t* are

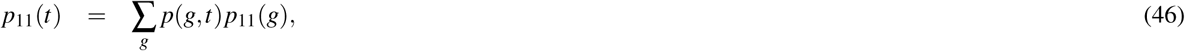

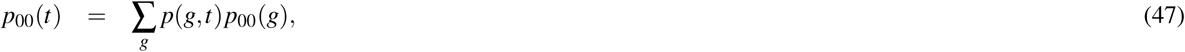

and the probability of observing an asymmetric division from a stem cell selected from a mouse of age *t* is *p*_01_(*t*) = 1 - *p*_11_(*t*) - *p*_00_(*t*).

#### Mathematical model of age-structured proliferation

The following model describes the evolution of stem cell numbers with age, taking into account differences in stem cell regenerative potential with number of times divided previously:

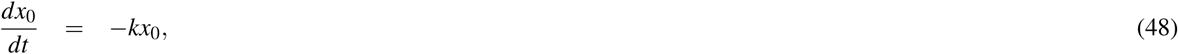

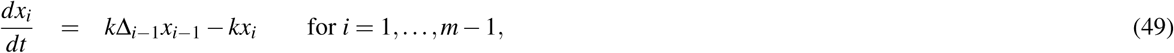

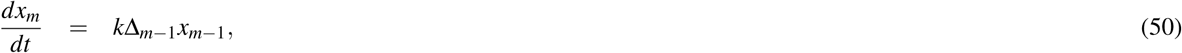

where *x*_*i*_(*t*) is the expected number of stem cells that have divided *i* = 0, 1, 2, 3, *…, m* times previously, Δ_*i*_ is the expected number of stem cells produced per division for a stem cell that has divided *i* times previously, and ln(2)*/k* is the expected cell cycle time. This model assumes that stem cells adopt a dormant state once they have divided *m* times. Experimentally we do not observe *x*_*i*_(*t*) but rather the total stem cell number Σ_*i*_*x*_*i*_(*t*), therefore it is model estimates of total cell numbers that are compared to experimental observations. We find that the best fit to the experimental data occurs when cells are allowed to divide four times before entering a state of dormancy (i.e. when *m* = 4). In particular, because our PDC experiments indicate that Δ_*i*_ → *-*1 for large *i* (i.e. five or more), adding additional divisions results in a loss of stem cell numbers with age which is not observed experimentally.

## Supplemental Figures

**Figure S1.**
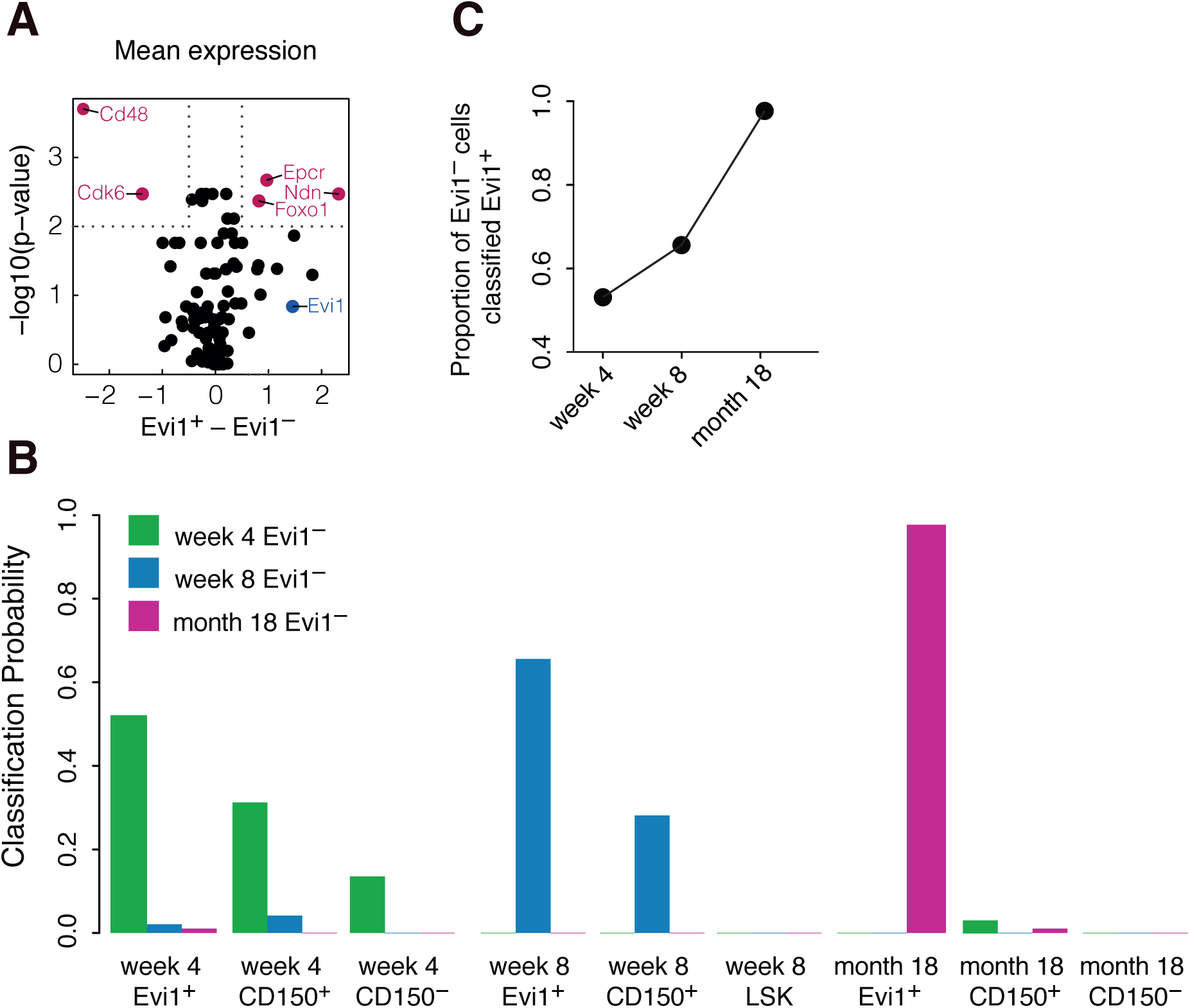
Single cell profiling of the Evi1^−^ fraction. (A) Volcano-plot of single cell gene expression differences between the Evi1^+^ and Evi1^−^ fractions (log-fold-change of mean expression) and negative log of p-values (FDR corrected using the Benjamni-Hochberg [BH] method) from a hypothesis test for single cell differential gene expression (McDavid et al. 2013). Significant genes (FDR corrected p-value < 0.01, and fold change > 1.4) are highlighted in cerise; Evi1 (highlighted in blue) is substantially upregulated (fold-change > 2) in the Evi1^+^ fraction, but does not pass the corrected significance threshold. (B) Results of classification of Evi1^−^ cells using our ANN classifier. (C) The proportion of Evi1^−^ cells classified as Evi1^+^ increases with age, indicating that these two populations become more alike with age.

**Figure S2.**
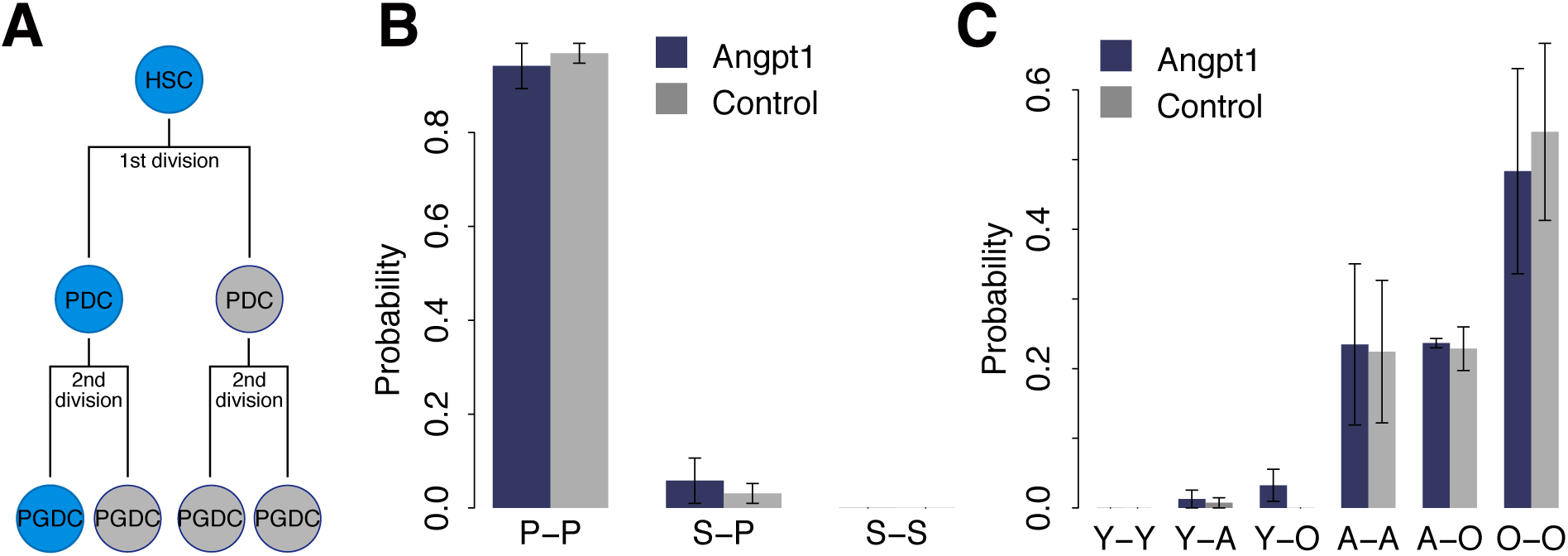
Comparison of paired granddaughter cells. (A) Schematics of paired granddaughter (PGDC) cell assay. (B) Classification of PGDC divisions using our trained ANN, restricted to regenerative status. Most secondary divisions in culture give rise to daughters with progenitor characteristics. (C) Classification of PGDC divisions using trained ANN, restricted to age. Spontaneous aging is very commonly observed in one or other, or both, granddaughter cells. Spontaneous rejuvenation is never observed, to within accuracy of classifier. In all panels data are expressed as the mean ± the standard deviation. Representative data from 3 independent biological replicates (with 48 pairs of division per replicate) are shown.

**Figure S3.**
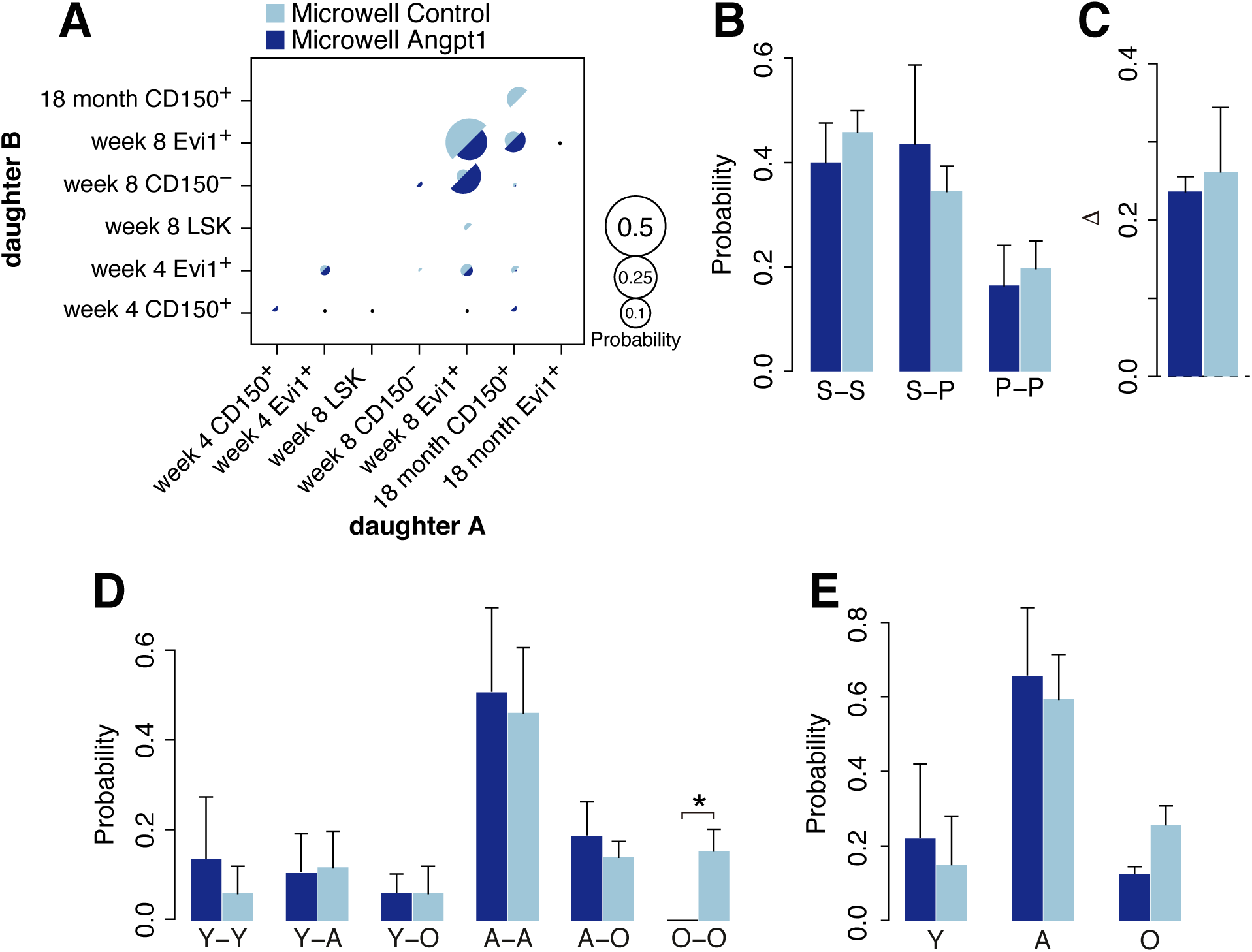
Division patterns of LT-HSCs cultured in PEG microwells. (A) Classification of divisions from adult (8-week old) stem cells cultured in PEG microwells using trained ANN. Daughter cell identities are not ordered. (B) Classification restricted to regenerative status. A range of divisions is observed in both control and Angpt1 treated conditions. (C) The expansion factor Δ is positive in both control and Angpt1 treated conditions indicating robust expansion of the stem cell pool. (D) Classification restricted to age. Both daughter cells from the majority of divisions retain appropriate age characteristics, however both spontaneous rejuvenation and aging are seen in a small proportion of cells. (E) Age distribution of daughter cells, without regard for pairing. In all panels data are expressed as the mean ± the standard deviation. Representative data from 3 independent biological replicates (with 48 pairs of division per replicate) are shown.

**Figure S4.**
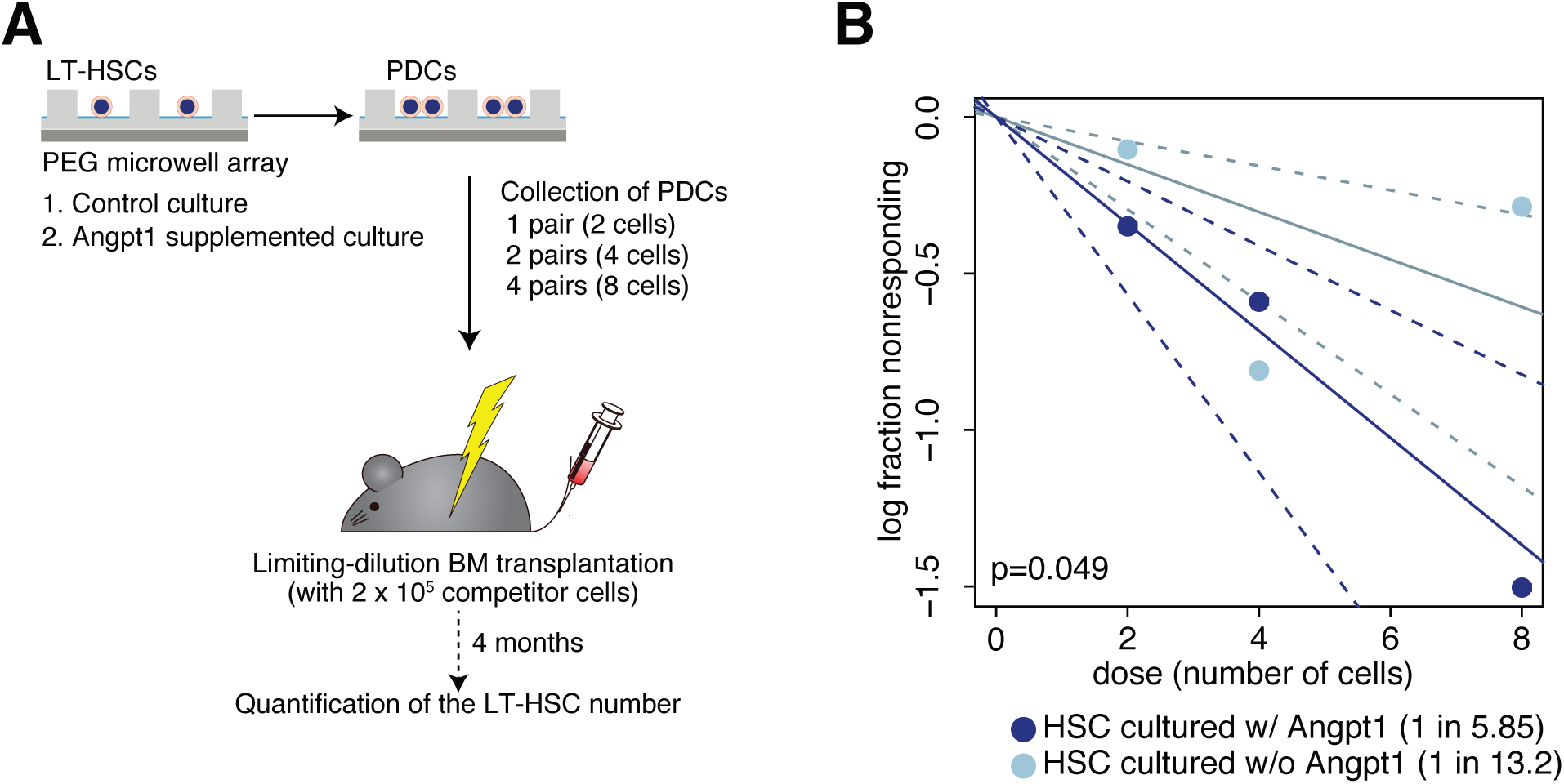
Microwell culture enhances LT-HSC self-renewal. (A) Schematic of limiting dilution bone marrow transplantation (BMT) assay. (B) Limiting dilution BMT of cultured PDCs shows that microwell culture maintains HSC activity *ex vivo* with and without Angpt1 treatment. In all panels data are expressed as the mean ± the standard deviation. Representative data from 3 independent biological replicates are shown. p-values determined by extreme value limiting dilution analysis (ELDA) in all panels (Hu & Smyth 2009).

**Figure S5.**
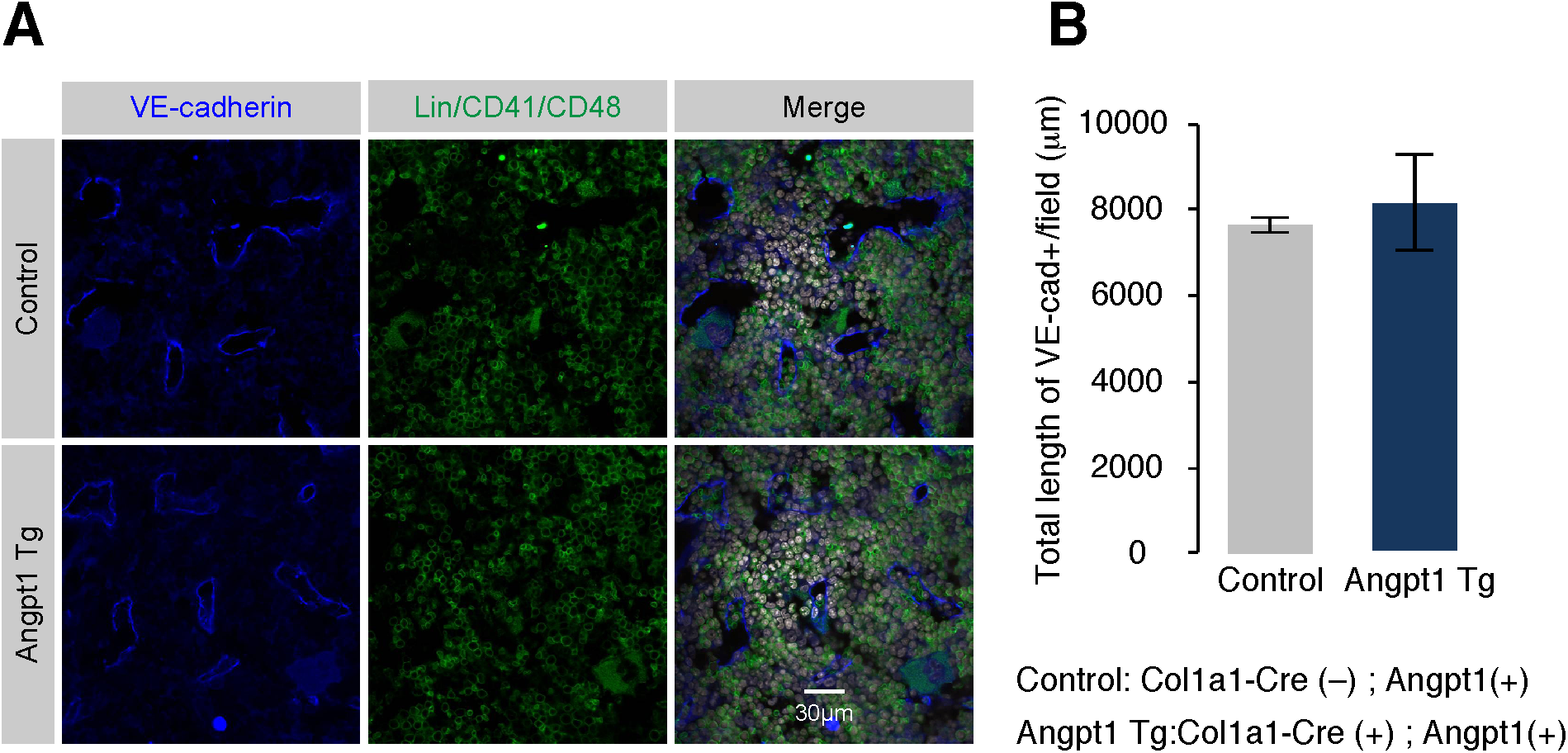
Comparison of vascular length in control and Angpt1 Transgenic mice. (A) Immunohistochemical staining of VE-cadherin (blue) and lineage markers/CD41/CD48 (green). Nuclei are stained with TOTO3 (white). Scale bar is 30*µ*m. (B) Total length of VE-cadherin^+^ vasculature per field. Data are expressed as the mean ± the standard deviation (control: 154 fields, Angpt1 Tg: 170 fields). Representative data from 3 independent experiments are shown.

**Figure S6.**
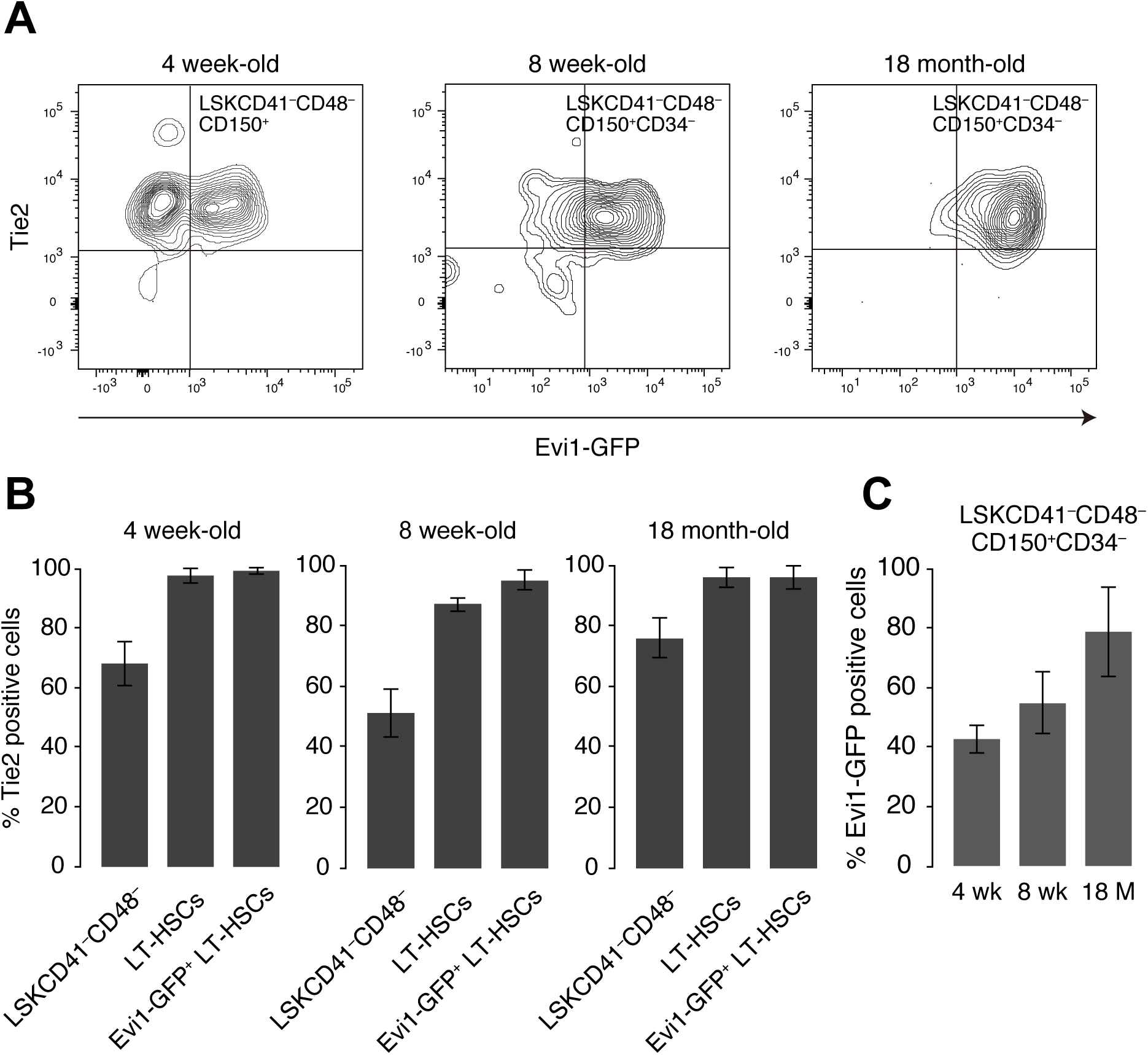
Angpt1 and Tie2 expression in young, adult and aged HSCs. (A) Coexpression patterns of Angpt1 and Tie2 throughout life. (B) LT-HSCs retain Tie2 expression with age. (C) The proportion of Evi1^+^ cells within the LSKCD41^−^CD48^−^CD150^+^CD34^−^ fraction increases with age. In all panels data are expressed as the mean ± the standard deviation. Representative data from 3 independent experiments are shown.

